# When does sexual selection through mate choice deplete versus exaggerate genetic variation: is there a lek paradox?

**DOI:** 10.1101/2024.12.10.627784

**Authors:** Kuangyi Xu, Maria R. Servedio

## Abstract

The evolution of female preferences for male display traits relies on females receiving indirect benefits from their mate. This requires substantial genetic variation in display traits or male quality. Nevertheless, sexual selection through mate choice has been assumed to deplete this genetic variation, ultimately diminishing strong preferences. However, sexual traits often have higher genetic variation than non-sexual traits. This contradiction, and, relatedly, how costly preferences and display traits are maintained, is called the “lek paradox”. Using infinitesimal models, we show that sexual selection through mate choice allows variation in male display traits and female preferences to mutually exaggerate each other, in a process analogous to runaway sexual selection but in terms of genetic variation. Therefore, contrary to prior suppositions, sexual selection may increase equilibrium genetic variance in display traits and preferences over that under random mating, provided that the variance of female preferences relative to male traits during mate choice is not too small and preferences are not too weak. Notably, even if equilibrium preference variation is substantially smaller than trait variation, trait variation can still increase over that under random mating, provided selection on the trait compared to the preference is sufficiently strong. Furthermore, when trait variation does decrease, this reduction is generally slight. It is under situations such as lekking, where females can simultaneously choose from many males so that mate discrimination is effectively strong and fitness costs of choice weak, that sexual selection through mate choice may be most powerful in exaggerating genetic variance in male displays.

**Significance statement:** The “lek paradox”—the dissonance between a hypothesized loss of variation in sexual display traits due to mate choice, leading to the subsequent cessation of sexual selection, and evidence of high variation in such traits in nature, accompanied by the long-term persistence of sexual selection—is an enduring mystery of the sexual selection literature. We clarify and quantify multiple pathways by which sexual selection via mate preferences alters genetic variance in both display traits and female preferences. Using mathematical models, we show the lek paradox is built upon a false premise. We find a wide range of conditions under which sexual selection increases or minimally reduces variation in display traits, allowing the maintenance of substantial variance in traits and preferences.

## Introduction

The evolution of exaggerated mating display traits in males and preferences for these seemingly non-adaptive traits in females have been questions of long-standing interest in evolutionary biology. Darwin (1871) proposed that elaborate traits in males may evolve because they increase the mating success of individuals that possess them. Later work elucidated that female preferences for male traits that offer no direct benefit can evolve because females may gain indirect genetic benefits. Specifically, females with a preference that mate disproportionately often with a male bearing a display trait tend to produce offspring carrying the alleles for both the attractive trait and the preference; this genetic correlation between preferences and traits will lead to their coevolution (the “Fisher process”; Fisher 1958). In addition to this process, females with a preference may produce more viable offspring, as occurs when males bearing more exaggerated traits have better genetic condition (e.g., a good genes mechanism: Jones & Ratterman 2009, Kuijper et al., 2012, Dhole et al. 2018). The coevolution of traits and preferences by these processes can theoretically cause their substantial exaggeration. However, theory suggests that for the indirect genetic benefits generated by sexual selection to be strong enough for exaggeration to occur, substantial genetic variation in male display traits and strong enough female preferences are required (Lande 1981, Kirkpatrick 1982, Iwasa et al. 1991, Kirkpatrick & Barton 1997, Hall et al. 2000, Jones 2009, Fry 2022, Xu et al. 2023, Servedio 2025).

However, it is unclear how genetic variation in male traits and persistent preferences can be maintained in a population. A long-standing claim in the literature is that sexual selection via female preferences should deplete genetic variation in male display traits (e.g., Borgia 1979, Kirkpatrick & Ryan 1991, Kotiaho et al. 2008). This depletion would reduce genetic benefits from mate discrimination, making females unable to evolutionarily maintain strong preferences if preferences incur fitness costs (Lande 1981, Kirkpatrick & Ryan 1991, Hall et al. 2000). As sexual selection weakens through this process, costly elaborated male traits will also be hard to maintain. Contrary to this expectation, exaggerated sexual traits and strong female preferences appear in many taxa (Andersson 1994, Rosenthal 2017, Prum 2018). Additionally, genetic variance in sexual traits is not unusually low; in fact, empirical evidence finds greater genetic variance and a similar level of heritability in sexual compared to non-sexual traits (Pomiankowski & Møller 1995, Prokuda & Roff 2014). This contradiction is known as the “lek paradox” (Borgia 1979, Kirkpatrick & Ryan 1991, Kotiaho et al. 2008), named after the “lek” breeding system in which females choose from an arena of displaying males that give no resources or other direct benefits. The term “lek paradox” has been used by some authors to refer primarily to the ongoing presence of strong preferences in the absence of direct benefits (e.g., Taylor & Williams 1982, Kirkpatrick & Ryan 1991), and by others to refer to the maintenance of genetic variation in male traits (e.g., Kotiaho et al. 2008), but as discussed above these issues are inextricably intertwined.

One possible way that trait variation may be maintained under mate choice is when variation in male trait values preferred by females is sufficiently larger than the actual variation in male traits. In this case, such as when there is high variance in unimodal preferences, these preferences may impose disruptive selection on male traits, increasing their variance (Lande 1981, van Doorn & Weissing 2002, Weissing et al. 2011). However, this mechanism is not often invoked as a solution to the lek paradox, perhaps because it is unclear how the prerequisite of sufficient variation in preferences would be achieved in the first place (Weissing et al. 2011). In fact, previous studies suggest that the runaway process is unlikely to result in sufficient preference variation to meet this condition (Higashi et al. 1999, van Doorn et al. 2004).

Several other hypotheses, involving more than just the coevolution of preferences and traits, have also been proposed to solve the lek paradox, but they generally remain untested (Miller & Moore 2007, Kotiaho et al. 2008). Most of these hypotheses focus on cases where the expression of sexual traits depends on an individual’s genetic condition, which is genetically complex and maintains large genetic variation (Miller and Moore, 2007, Petrie 2021). Rowe and Houle (1996), for example, propose that high mutation rates on condition, which is determined by a large number of genes, may increase the genetic variance in sexually selected traits through the process of ‘genic capture’ (but see Kirkpatrick 1996). Nevertheless, empirical tests suggest that condition-dependent expression may not be sufficient to maintain genetic variance in sexually selected traits (Van Homrigh et al. 2007).

Despite long-standing efforts to solve the lek paradox, the central conceptual hypothesis that sexual selection through mate choice should deplete genetic variance in male traits has not been rigorously examined. Here, we use the infinitesimal model framework (Barton et al. 2017) in a “proof of concept” model (Servedio et al. 2014) to investigate the evolution of genetic (co)variances of preferences and traits under the Fisher process and a good genes mechanism. We find that rather than depleting genetic variation in male traits, sexual selection through mate choice can cause the genetic variation in expressed display traits and female preferences to mutually exaggerate each other when compared to the case of random mating. This result depends on two conditions: 1) variation in female preferences relative to the male trait cannot be too low (although it can, in some cases, be lower than trait variation), and 2) preferences cannot be too weak. In other cases, we find that trait variation may indeed be reduced by sexual selection, but this reduction is generally minimal, far from the loss of variation proposed by the lek paradox. As part of a thorough categorization of the sources of change in variance, we find that one, but not the only, key source of these results is that sexual selection through mate choice generates assortative pairing of males and females based on the display trait genes that they carry and/or express, thereby increasing the between-family variance among offspring (Falconer & Mackay, 1996). The lack of recognition of this mechanism may have led to an overestimation of both the minimum female preference variance required for mate choice to increase trait variance and the reduction of trait variance by sexual selection. Our analysis suggests sexual traits may often have larger genetic variation than non-sexual traits, as found in nature (Pomiankowski & Møller 1995, Bakker 1999), providing a reconciliation between the theoretical expectations and empirical patterns.

## Results

We consider a diploid, polygynous population with non-overlapping generations to investigate two mechanisms of sexual selection. First, in a model of the Fisher process, we consider two sex-limited polygenic traits: females with preference *y* choose males based on their display trait phenotype *x*. Second, we investigate the handicap model by including an additional trait *z*, associated with general viability and expressed in both sexes. Here females choose males based on the realized male trait *s* = *x* + *z*; *s* is thus an honest signal of male quality *z* (Iwasa et al. 1991). In both scenarios, all traits are polygenic with phenotypic values being normally distributed.

Each generation first undergoes viability selection on trait *z* in both sexes (only in the handicap model), on the display trait *x* in males, and on the preference *y* in females, which is assumed to be costly (with greater costs paid when the males that females prefer are rarer). Viability selection is followed by sexual selection via mate choice, by reproduction, and finally by mutation. We adopt the infinitesimal model, which assumes that the genetic value of offspring within a family is normally distributed around the mean of the parents’ trait values, with the variance within a family being independent of the parental trait values and not being affected by selection. These assumptions hold well when trait values are determined by the sum of many small-effect loci (Barton et al. 2017, 2023; Parsons & Ralph 2024). Viability and sexual selection change the distribution of genetic values among mated pairs, thereby affecting the evolution of genetic (co)variances. Details of the model and the recursions for the genetic variances are described in *Materials and Methods* and the SI Appendix.

The probability that a female with preference *y* will mate with a male with phenotype *x* or *s* is described by the preference function. For the Fisher process, we consider the three preference functions *ψ*(*x*|*y*) used in Lande (1981): 1) absolute preferences where 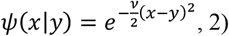 relative preferences where 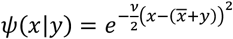, and 3) psychophysical preferences where *ψ*(*x*|*y*) = *e*^*xy*^. Here a higher *v*, which describes the preference strength, yields a stronger preference. Absolute and relative preferences impose stabilizing sexual selection, whereby females with preference *y* are most likely to mate with males with phenotype *x* = *y* or 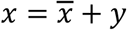, respectively. In contrast, under psychophysical preferences, females prefer males with more extreme phenotypes so that sexual selection is directional, and *y* denotes both the direction and strength of preference.

For the handicap model, the male trait *x* is replaced by the realized male trait *s* in the preference functions. Since traits *x* and *z* interact additively to comprise trait *s*, under absolute or relative preferences, stabilizing sexual selection often generates a negative genetic association between *x* and *z*. Under psychophysical preferences, directional sexual selection instead generates a positive genetic association between *x* and *z*.

### The Fisher Process

The recursion for the genetic variance of the male trait 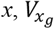, can be written as sum of contributions from several sources (see *Materials and Methods*), as

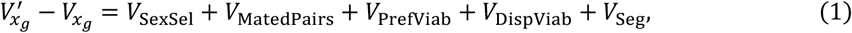

where 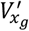 is the genetic variance in the next generation. The terms *V*_SexSel_ and *V*_MatedPairs_ are key to understanding the effects of mate choice on the change in trait variance. As described below and in Table 1, they can either in combination or, in the case of *V*_MatedPairs_, independently account for why trait variance may often be higher when there is mate choice versus when there is random mating.

**Table 1.**
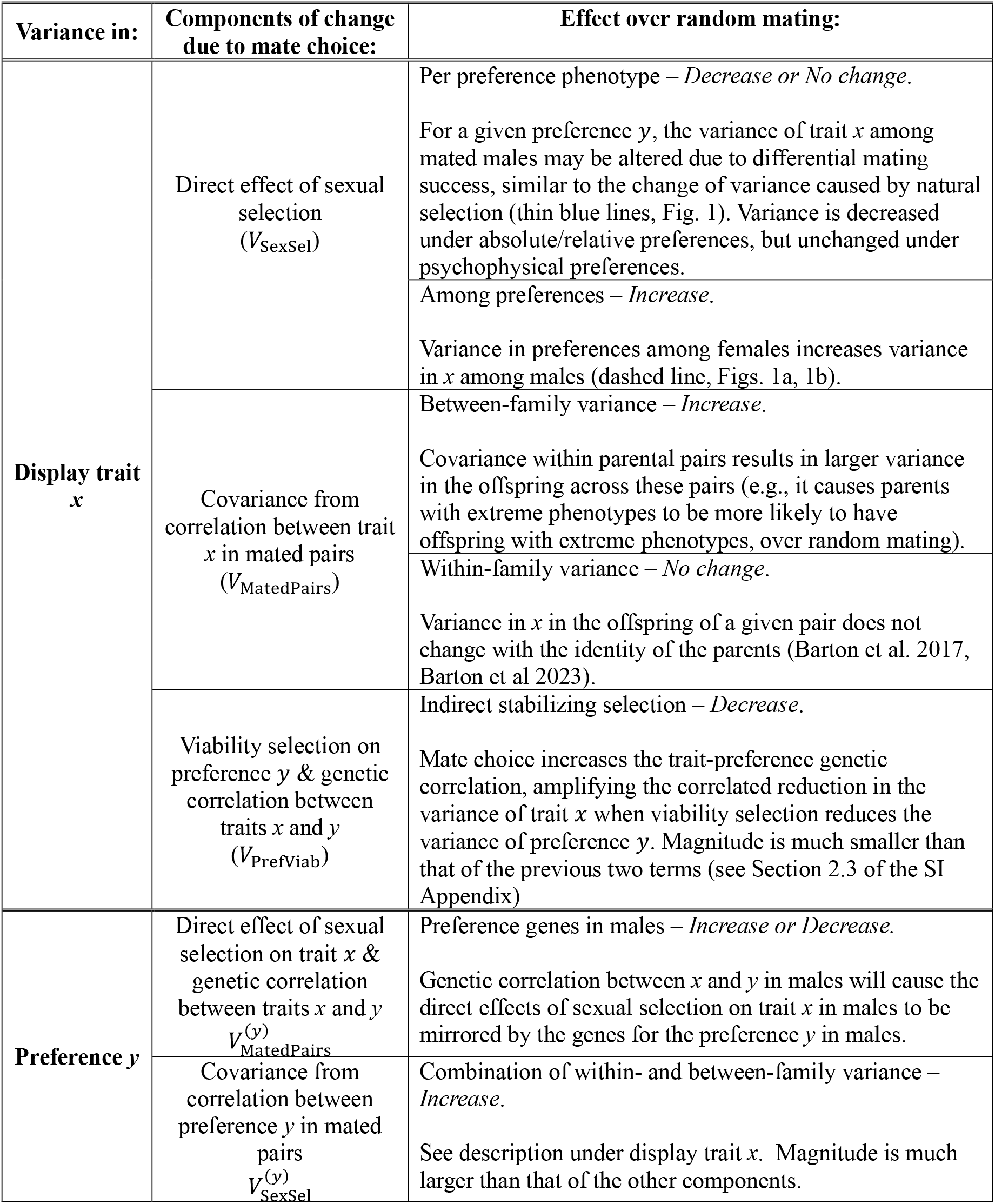
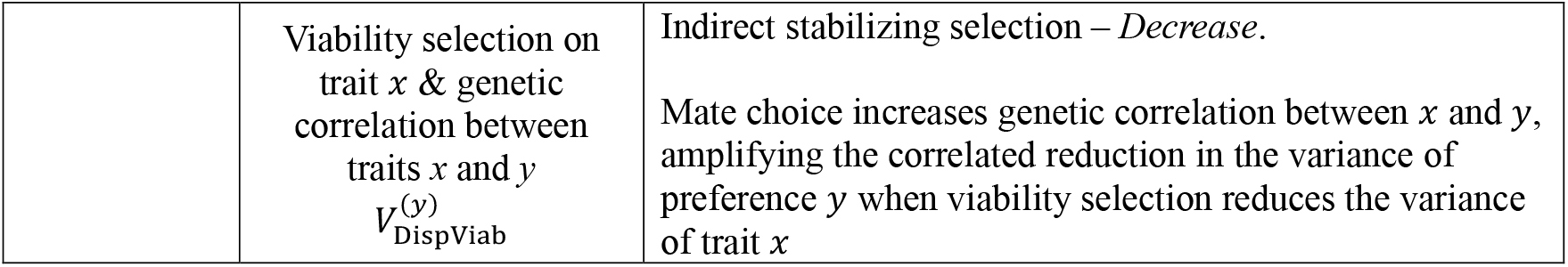
Components of change in the variance in display trait and preference due to mate choice.

The first term in Equation (1), *V*_SexSel_, describes the change in the variance of the male trait that arises from the direct effect of sexual selection as a selective force. Provided that preference functions are relative or absolute, and thus sexual selection is stabilizing, the variance in trait *x* among the successfully mated males chosen by the set of females with a specific preference phenotype *y* will be lower than the variance of trait *x* in the population at large (thin blue lines in Figure 1a). However, among females in the population there is a distribution of preference phenotypes. When summed across all values of *y*, the individual distributions (per preference *y*) of successfully mated male trait phenotypes will combine to result in a greater variance of successful male traits in the population as a whole. Indeed, this can result in a variance among successfully mated males that is higher than that in the display trait *x* before mate choice (Figure 1a). Specifically, we find that the direct effects of sexual selection will lead to an increase in male trait variance (*V*_SexSel_ > 0) when 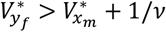, a condition that is more likely to hold when preference strength *v* is stronger, where 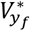 is the variance in preference *y* in females after viability selection and 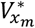 is the variance of *x* in males after viability selection (Section 2.1 of the SI Appendix). This is an extension of the condition identified under the limit of weak selection in the approximation by Lande (1981).

**Figure 1.**
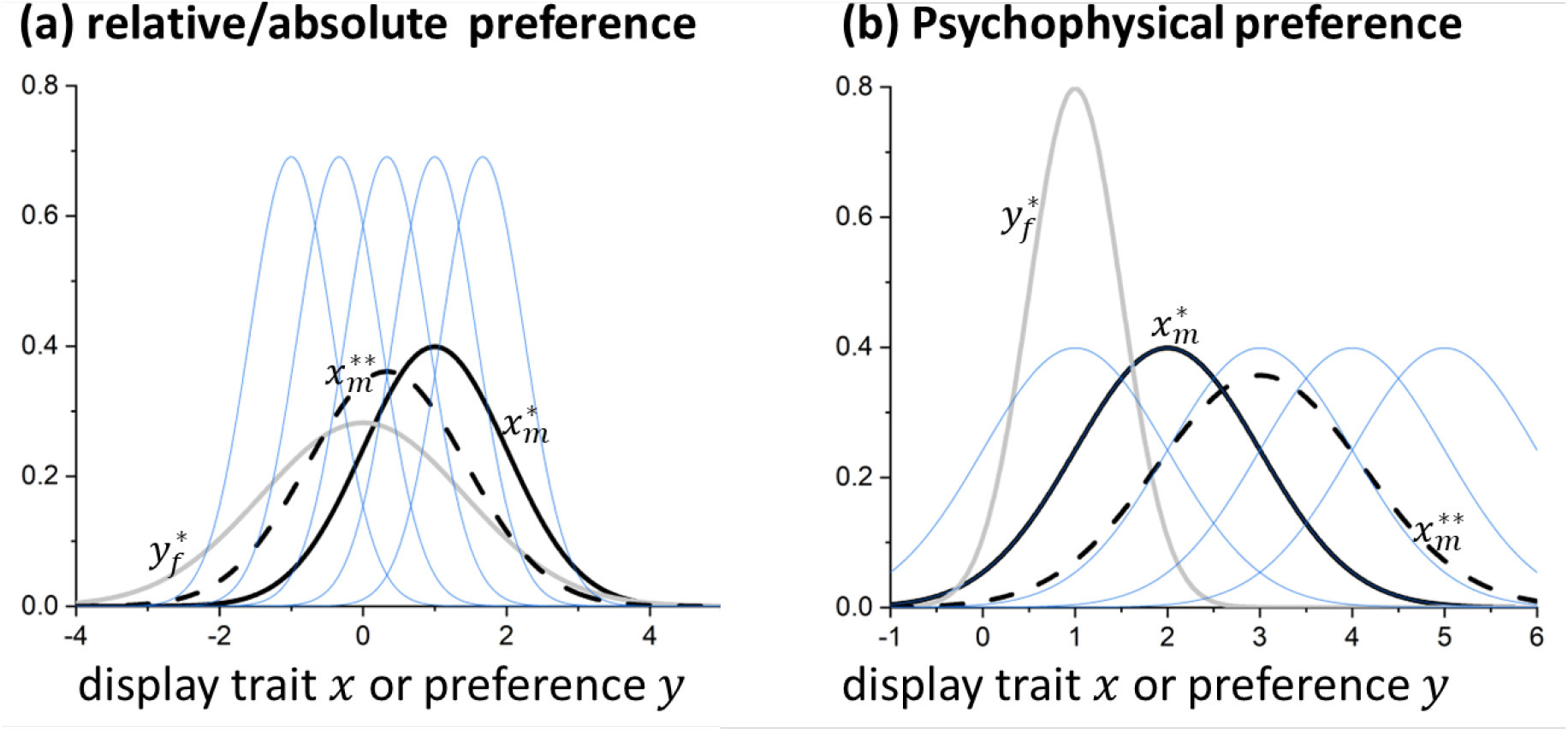
Direct effect of sexual selection through mate choice on changing the variance of display trait *x* in males. Black lines show the distribution of trait *x* in males before mate choice (solid) and in successfully mated males after mate choice (dashed). The thick grey line shows the distribution of preference *y* among choosing females. Each thin blue line shows the distribution of trait *x* among males mated with females with a given preference *y* (derived in Section 2.1 of the SI Appendix). (A): relative/absolute preferences, with 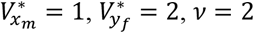. (B): psychophysical preferences, with 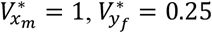.

When the preference function is psychophysical, the variance in trait *x* among successfully mated males per female preference value *y* remains the same as that before mate choice (thin blue lines in Figure 1b). The variance in trait *x* therefore uniformly increases through *V*_SexSel_ with psychophysical preferences provided there is some variation in female preference values (dashed line in Figure 1b). However, this result is not general to all open-ended preferences but is due to the special form of the psychophysical preference function (see Section 4 of the SI Appendix).

Unlike *V*_SexSel_, *V*_MatedPairs_, the variance resulting from the correlation in the values of trait *x* between mated pairs, always causes an increase in the variance of *x* compared to random mating, under which this variance is absent (with random mating *V*_MatedPairs_ = 0). Specifically, non-random mate choice generates positive covariance between females with preference *y* and males with display trait *x* in mated pairs. Preference and display trait genes also become correlated within individuals in the population, as offspring of both sexes inherit preference genes from their mother and trait genes from their father. Mate choice therefore generates covariance between trait *x* in mothers and trait *x* in fathers. In other words, mate choice is assortative with respect to the display trait genes (and also the preference genes). A mother carrying trait genes at the extreme of the distribution of *x*, for example, will be more likely to obtain a mate at the same extreme, over the case of random mating. She will consequently be more likely to have offspring with similarly extreme trait values. This effect of assortative mating by trait value, captured by *V*_MatedPairs_, therefore increases the between-family variance in trait *x* in the offspring generation over the case of random mating (Section 2.2 of the SI Appendix). Larger variation in the male display trait or female preference during mate choice increases the contribution of *V*_MatedPairs_, since *V*_MatedPairs_ is proportional to the trait-preference correlation within individuals, which is higher with larger variance of display trait and preference (Hall et al. 2000).

On the other hand, although mate choice increases the similarity in genetic values between paired individuals, for traits controlled by many loci, the variance among siblings (within-family variance) should not change significantly between random versus non-random mating. Intuitively, the genetic variation among siblings is mainly generated by genetic processes such as segregation and recombination during meiosis, represented by the term *V*_Seg_ in Equation (1). This variation mainly depends on the genetic relatedness between parents (Barton et al. 2017, 2023), but assortative mating only slightly elevates this relatedness when traits have a polygenic basis (Crow & Felsenstein 1982). Therefore, an overall increase in variance through *V*_MatedPairs_ should generally apply when traits are controlled by many loci.

The third term in Equation (1), *V*_PrefViab_, describes the decrease in the variance of trait *x* due to viability selection on preference *y* in females and the genetic correlation within individuals between *y* and *x*. Mate choice may increase the magnitude of this decrease over random mating by increasing the genetic correlation. However, the effect of mate choice through *V*_PrefViab_ tends to be much smaller than that through either *V*_SexSel_ or *V*_MatedPairs_ (Figure S2; see Section 2.3 of the SI Appendix for explanations) and thus is not our focus.

The last two terms in Equation (1) are not expected to change across the cases of random mating and mate choice. V_DispViab_, which is negative, is the genetic variance that is reduced by stabilizing viability selection on trait *x*. Finally, *V*_Seg_, the variance due to segregation, mutation and recombination—often referred to as segregation variance—is largely insensitive to mate choice, as explained above.

Therefore, the net effect of mate choice over random mating on the equilibrium genetic variance, which is reached rapidly (Figure S1), mainly relies on whether any potential decrease in variance that may occur through *V*_SexSel_ can be overcome by the increase over random mating that necessarily occurs through *V*_MatedPairs_ (Figure S2). Under psychophysical preferences, since *V*_SexSel_ is positive, mate choice always leads to larger genetic variances of traits and preferences than random mating (Figures S6, S7). Under absolute or relative preferences, the effects of mate choice on the variances are less clear, as *V*_SexSel_ can be negative; this is therefore our focus below. We illustrate the conditions under which equilibrium genetic variances under mate choice are larger or smaller than those under random mating. To gain mechanistic intuition of these results, we also derive expressions for the conditions when stronger preference strength increases the variance after one generation, by considering the case where there is no environmental variance (see Section 2 of the SI Appendix).

Our equilibrium analysis shows that whether, through the multiple effects above, mate choice will overall increase or reduce the equilibrium male trait variance over random mating depends largely on the preference strength, as well as the magnitude of variation in male traits and preferences, which are, in turn, predominantly determined by segregation variance and the strength of viability selection. Importantly, mate choice can often increase the equilibrium trait variance over random mating. In Figures 2a and 2d, when the preference strength is sufficiently large (y-axis values above the white dashed line), the equilibrium trait variance exceeds that under random mating (which is approximately equal to the values seen on Fig. 2 when the preference strength *v*=10^−3^). This increase over the case of random mating can occur even when the equilibrium preference variance, 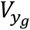, is much smaller than the trait variance (compare Fig. 2d with 2e), provided that viability selection on the display trait and the preference strength are sufficiently strong (the region above the white dashed line in Figure 2d).

**Figure 2.**
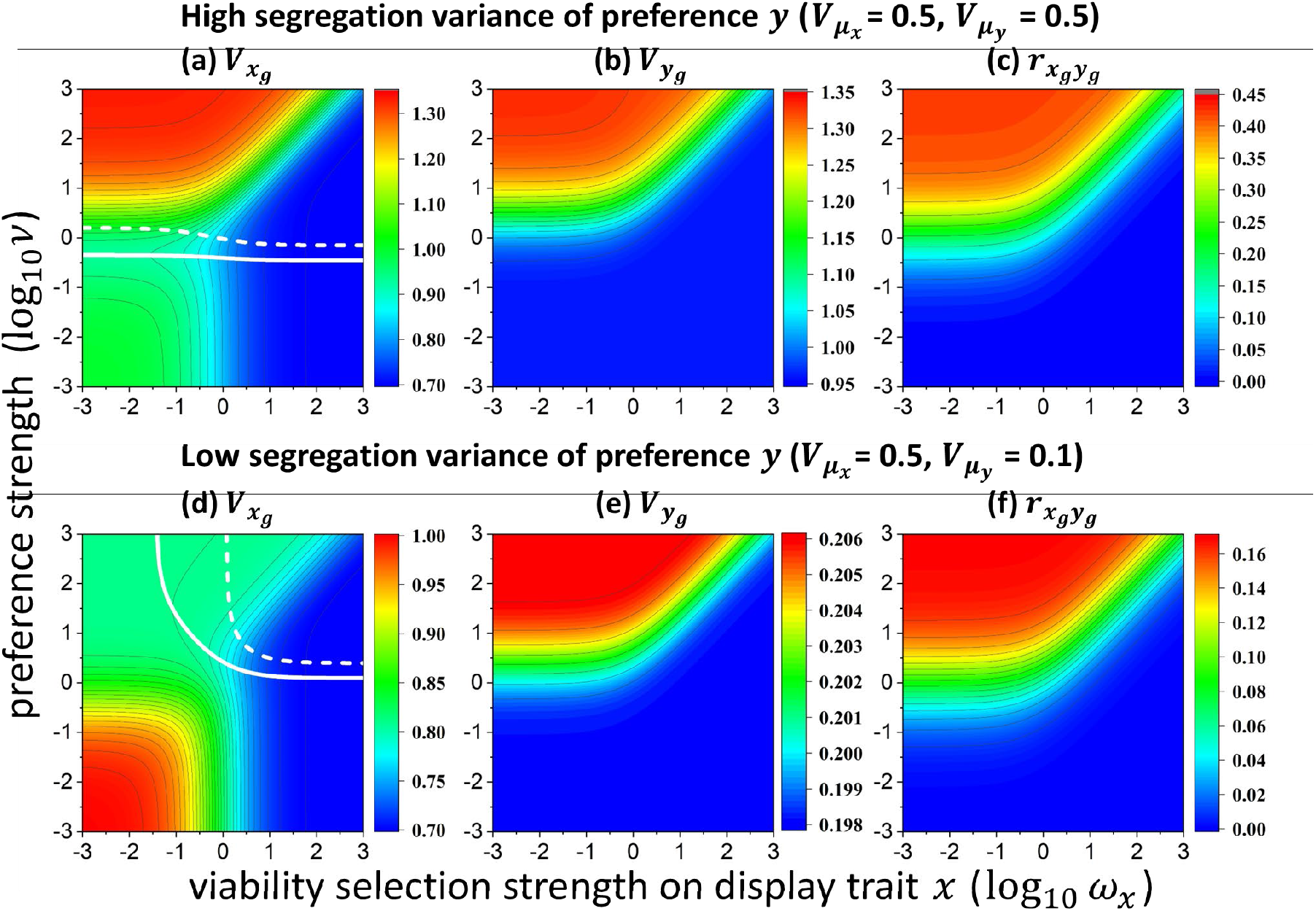
Equilibrium genetic variance of the male trait *x* and female preference *y*, and the genetic correlation between the two traits under absolute or relative preferences in the Fisher process model. In in panels (a) and (d), the solid white line marks the preference strength that minimizes the genetic variance of trait *x*, and the dashed white line marks the preference strength above which mate choice increases trait variance over random mating. In panel (d), the slopes of both lines converge to infinity at specific values on the X-axis. Below the X-axis value corresponding to the white solid line, stronger preference monotonically reduces trait variance; below the value for the white dashed line, mate choice cannot increase trait variance beyond that observed under random mating, even when the effect of preference strength is non-monotonic. Results are obtained from numerical iteration of Equation (3). Other parameters: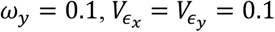. Note the differences in the color scale for each panel.

Specifically, when the variance of the preference compared to the male trait is not too small, the equilibrium genetic variance of trait 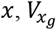, first decreases and then increases as preference strength becomes stronger, and therefore is minimized at an intermediate preference strength (solid white line in Figures 2a, 2d). In turn, this case occurs when the segregation variance of the preference is not too small compared to that of the male trait (Figures 2a, S2), or when the segregation variance of the preference is small but viability selection on the male trait is strong enough (region spanned by the white solid line, Figure 2d).

These non-monotonic changes can be explained as follows. We have learned that *V*_SexSel_ is determined by a combination of the reduction of display trait variance by mate choice per preference phenotype, and the increase due to variation in preference values among females (Figure 1a). When the preference strength is below a critical level 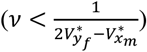, increasing the preference strength elevates the magnitude of both effects but has a stronger effect on the former (see Section 2.1 of the SI Appendix), thereby reducing the genetic variance of trait *x*. Additionally, under weak preferences, the contribution from the correlation of *x* values between mated pairs, *V*_MatedPairs_, is weak due to a low trait-preference correlation (Figure S2), and thus cannot compensate for the loss of trait variance.

As preferences become sufficiently strong, an increase in preference strength will increase *V*_SexSel_ by more effectively elevating the positive contribution from the variation in preference values, potentially making *V*_SexSel_ positive (Figure S2; Section 2.1 of the SI Appendix). More importantly, stronger preferences also elevate the trait-preference genetic correlation (Figure 2c), causing the positive contribution from V_MatedPairs_ to outweigh the (potential) reduction caused by *V*_SexSel_ (Figure S2). Consequently, stronger preferences will increase the genetic variance of trait *x*, and can make it larger than that under random mating when preference strength exceeds a critical value (white dashed line in Figures 2a, 2d). This critical value, as plotted in Figure 3a, is lower as the ratio of viability selection strength on preference *y* to that on display trait *x* becomes smaller and the segregation variance of preference becomes larger (compare the red and black lines in Figure 3a). Also, given the same relative strength of viability selection on the preference and display trait, the likelihood for mate choice to increase the trait variance over random mating depends on the absolute magnitude of selection. A smaller magnitude of selection increases this likelihood when selection on the preference is stronger than on the display trait, or when selection on display trait is stronger but the segregation variance of the preference is not too low (compare the solid and dashed lines in Figure 3a). This is because the variation in the trait and preference is greater under weaker viability selection, which in turn increases the trait– preference correlation (Hall et al. 2000), and hence the positive contribution from *V*_MatedPairs_.

**Figure 3.**
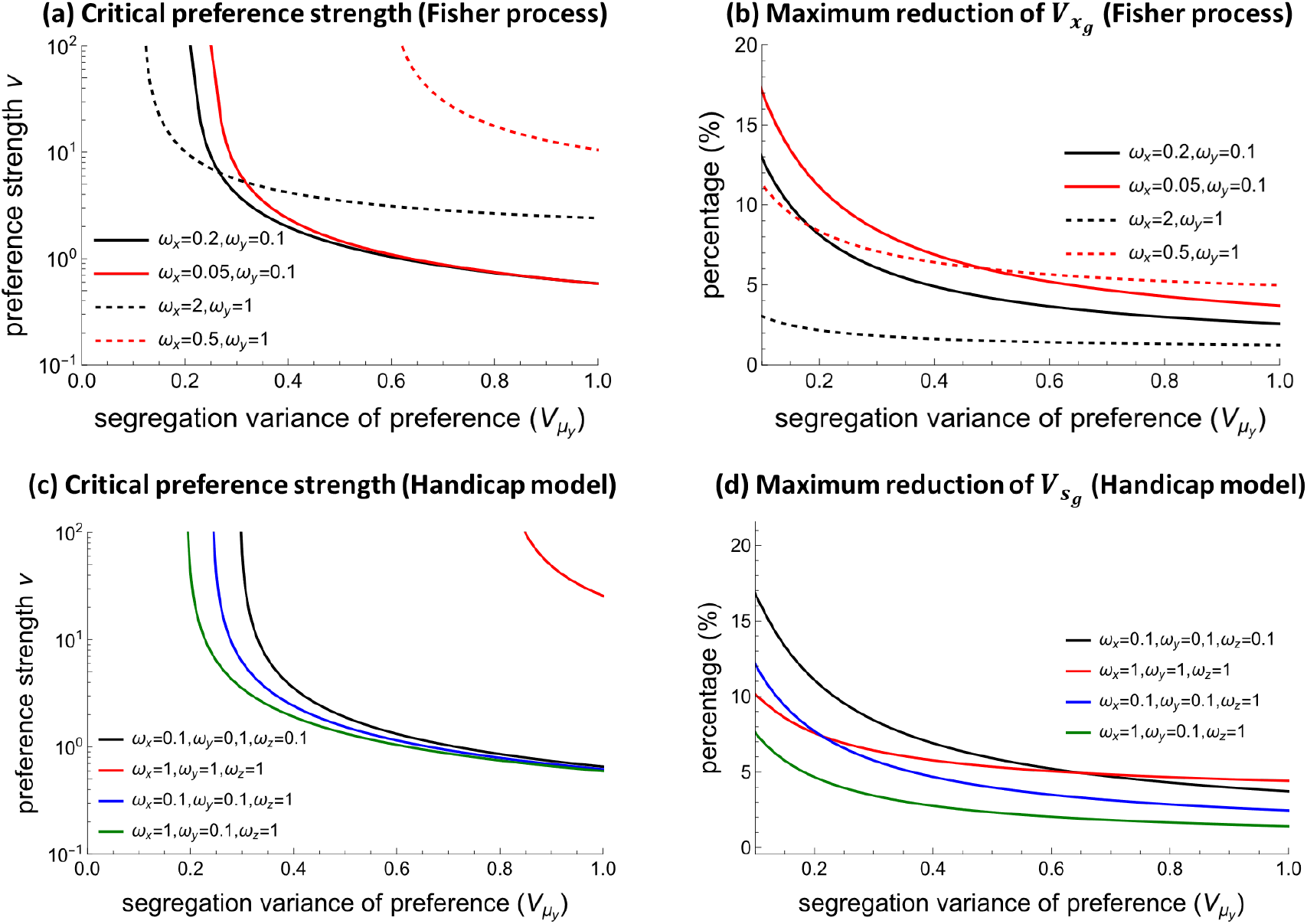
Panels (a) and (c): critical preference strength above which mate choice will lead to a larger equilibrium genetic variance of trait *x* than that obtained under random mating, with absolute or relative preferences (Fisher process, panel (a)) or realized display trait *s* (handicap model, panel (c)). Panels (b) and (d): maximum possible reduction of the equilibrium genetic variance of trait *x* (Fisher process, panel (b)) or trait *s* (handicap model, panel (d)) caused by mate choice with absolute or relative preferences relative to the variance under random mating. Results are obtained from numerical simulations. The segregation variance of trait *x* is 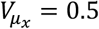 for the Fisher process, and 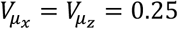 are used for the handicap model so that the magnitude of the variance of trait *s* is similar to that of trait *x* in the Fisher process. Other parameters are 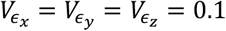.

However, stronger preferences will monotonically reduce the male trait variance when the variance of female preference is excessively small compared to the male trait. This occurs when the segregation variance of preference 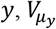, is relatively small and viability selection on the male trait is weak (left-hand side, Figure 2d), or when selection on preference is strong (Figure S3). Under these conditions, the direct effect of mate choice, *V*_SexSel_, is negative and often the dominant factor (Figure S2), since the trait-preference correlation, and hence *V*_MatedPairs_, remains weak (Figure 2f).

As can therefore be seen, there are cases when mate choice reduces trait variation over random mating, and this reduction either monotonically becomes larger with stronger preferences or reaches a maximum at intermediate preference strength (Figures 2a, 2d). However, in contrast with language commonly used in presentation of the paradox of the lek, the maximum possible reduction compared to the variance under random mating (e.g., where the equilibrium variance is at its minimum, along the white lines in Figure 2) is usually small, unless the segregation variance of preference relative to display trait is very low (Figure 3b).

Mate choice affects the genetic variance of the preference *y* through a recursion equation with components analogous to those in Equation (1) (see Table 1; Section 1.3 of the SI Appendix). Because there is a genetic correlation between trait *x* and *y* in males, the analog to 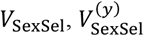, where the superscript indicates variance is in the preference, leads to a decrease or increase in variance of *y* in males compared to its value if mating were random, depending on whether variance in trait *x* increases or decreases due to *V*_SexSel_. Additionally, mate choice can increase the magnitude of the correlated reduction in the variance of *y* when viability selection reduces the variance of trait *x* in males, by increasing the correlation between trait *x* and *y* (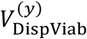, see Table 1). Finally, the covariance of values of *y* in paired females and males generated by mate choice, 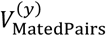, will lead to an increase of variance of *y* over random mating, where this covariance is absent (as described for the variance in *x* above).

Interestingly, when these three components are combined, mate choice always leads to a larger equilibrium genetic variance of preference *y* over the case of random mating. Stronger preferences likewise always increase the variance in *y*, regardless of the strength of viability selection on the preference and trait (Figures 2b, 2e). This is because the positive contribution through 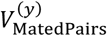 is always the dominant force (Figure S2). Intuitively, the change in the variance of *y* through 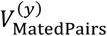 is proportional to the genetic correlation between traits *x* and *y*, while the changes through 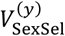 and 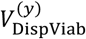 are proportional to the square of the correlation, and hence a weaker effect (see Section 5 of the SI Appendix for further explanation). Therefore, mate choice itself, by exaggerating variance in preferences, can partly prevent the potential decrease in variance of male trait due to mate choice.

### The Handicap Principle

When sexual selection by mate choice operates under the handicap principle, the results are similar to those found under the Fisher process. Importantly, we find that mate choice can increase the variance in condition (trait *z*) as well as that in the display (trait *x*). Specifically, under absolute or relative preferences, whether mate choice increases or reduces the variance of realized display trait *s* = *x* + *z*, and thus also the variance of its components *x* and *z*, over random mating depends largely on the relative magnitude of the variances in preference *y* and trait *s* at the stage of mate choice (this evolves to become higher under a larger segregation variance of trait *y* and stronger viability selection on trait *x*; compare Figures 4a vs 4e for trait *s*, and 4b vs 4f for trait *x*; also see Figure S4 for the effects of selection strength on preferences).

**Figure 4.**
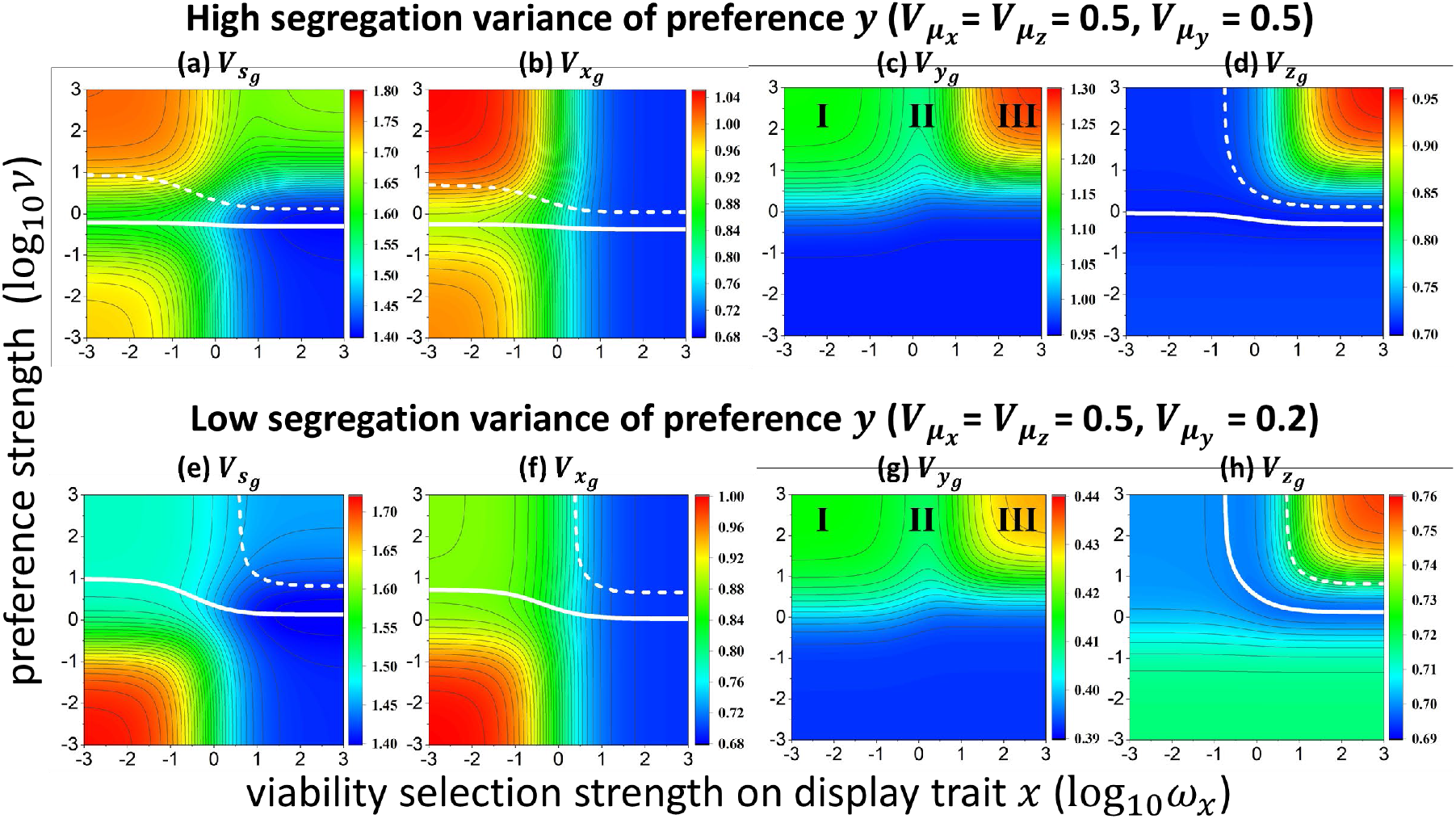
Equilibrium genetic variance of realized display trait *s* = *x* + *z*, display trait *x*, preference *y*, and viability-associated trait *z* under absolute or relative preferences for the handicap model. The biological meanings of white solid and dashed lines are the same as those in Figure 2. The genetic variance of trait *s* is calculated as 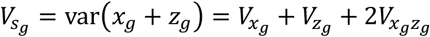. Other parameters are 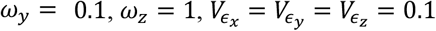.

Again, mate choice can lead to a larger variance in trait *s* (thus also traits *x* and *z*) than random mating when preference strength exceeds a critical value, which declines with larger segregation variance of the preference. Generally, mate choice is more likely to exaggerate trait variance over random mating when viability selection on the preference relative to the male trait is weaker, the absolute magnitude of selection on both trait and preference is weaker, and selection strength on the general viability trait *z* is larger (Figure 3c). The maximum possible reduction of the variance of trait *s* caused by sexual selection again tends to be small (Figure 3d). With psychophysical preferences, trait variance once again always increases over the case of random mating (Section 4 of the SI Appendix).

Stronger sexual selection always exaggerates the variance of female preference *y* under the handicap principle (Figures 4c, 4g), as under the Fisher process. Interestingly, this exaggeration is greatest when either trait *x* or *z* contributes a large proportion of the variance in the realized display trait *s* (regions I, III in Figures 4c, 4g, compare with Figure 4b, 4d, 4f, 4h), but becomes weak when traits *x* and *z* contribute comparably to the variance of trait *s* (region II in Figures 4c, 4g). Intuitively, in the former case, there is a strong genetic correlation between preference *y* with either trait *x* or *z*, but in the latter case, the genetic correlations between *y* and both traits *x* and *z* are weak (Figure S5).

The intuition above also explains the result that sexual selection is unlikely to have significant impacts on the genetic variance of both traits *x* and *z* simultaneously. Specifically, mate choice is effective in altering the variance of trait *x* or *z* when one of the two traits accounts for a large proportion of the variance of realized display trait *s*, but ineffective in affecting the genetic variance of the two traits when both traits contribute comparably (see Figures 4b, 4f for trait *x* and Figures 4d, 4h for trait *z*; Section 3 of the SI Appendix). This result is similar to the finding that the evolution of a preference for one display trait will block the evolution of preference for another display trait (Iwasa & Pomiankowski 1994). However, the mutual inhibition in the evolution of the variance of traits *x* and *z* may result from our assumption of an additive interaction of trait *x* and trait *z* on the realized trait *s*, and may be more complicated if this interaction instead takes a non-linear form.

## Discussion

The puzzle of why females should discriminate among males that offer no direct benefits has caused a flurry of speculation from as early as the 1950s (e.g., Shepard 1953, Fisher 1958, Lack 1968, Petit & Ehrman 1969, Trivers 1972), long before Borgia coined the term “The Lek Paradox”, popularizing this purported conundrum in 1979. At the heart of the lek paradox is the idea that the erosion of variance in male traits and/or condition due to viability and sexual selection will prevent females from reaping genetic benefits by mating with their preferred male (Borgia 1979, Kotiaho et al. 2008). Many theoretical and empirical studies have attempted to explain how variation in traits might be maintained, taking it as a given that variation will otherwise be eroded; a search on Google Scholar yields at least 14 papers with some variant of “resolution of the lek paradox” in the title alone. Solutions supported by mathematical models include preferences for heterozygosity (Fromhage et al. 2009, Aparicio 2011), fluctuations in selection on showy displays due to predator-prey cycles (Lerch & Servedio 2023), and the capture of non-additive genetic variation (Neff & Pitcher 2008) and of indirect genetic effects (Miller & Moore 2007; reviewed in Kotiaho et al. 2008). These solutions, and others in the literature, could indeed contribute to an increase in trait variation during the process of sexual selection. Additionally, we propose that there will in fact be no lek paradox under many conditions. Specifically, our models with evolving (co)variances of preferences and traits reveal that, as with the runaway process for mean trait values, selection can drive variation in traits and preferences to mutually increase one another, thereby often increasing or only minimally reducing genetic variation in male traits, while always exaggerating variation in preferences.

Stabilizing and directional viability selection generally deplete genetic variation, and the assumption at the heart of the lek paradox is that sexual selection will behave in a similar way. But in terms of its effect on genetic variances, sexual selection via mating preferences differs from viability selection in two primary ways, both of which can cause variance in display traits to increase. First, although matings from females with a specific preference value will reduce the variance of display traits within successfully mated males in a similar way as would viability selection, there is an additional contribution to display trait variance that arises from the fact that there is variation in preferences among females (Figure 1; see also Lande 1981; van Doorn et al. 2004; Weissing et al. 2011). This means that, overall, the variance in displays among successfully mated males may directly increase due to mate choice.

More importantly, we have identified another critical effect of mate choice on trait variance — the genetic value of the display trait will be positively associated between paired males and females (see *V*_MatedPairs_ in Table 1). This assortative mating with respect to the display trait will lead to an increase in the variance of offspring across all mated pairs (between-family variance), compared to the case of random mating. Together, these factors can cause an increase in display trait variance, over random mating, due to sexual selection by mate choice—directly contradicting the premise of the lek paradox.

In general, whether trait variance evolves to be larger or smaller under mate choice depends on the preference strength, as well as the relative magnitude of the variances in the preference and male trait. These are evolving variances, that are in turn affected by the relative magnitude of segregation variances and viability selection strength. Importantly, and contrary to previous findings (Lande 1981, Roff & Fairbairn 2014), when viability selection on female preference is weak compared to on the display trait, mate choice can increase the display trait variance even when the segregation variation of the preference is substantially smaller than that of the display trait, provided preferences strength is sufficiently strong (Figure 3a). This may help to explain why sexually selected traits can exhibit higher genetic variation than non-sexual traits (Pomiankowski & Møller 1995, Bakker 1999). On the other hand, display trait variance does evolve to be smaller under mate choice when preferences are weak or variation in the preference compared to the display trait remains small during coevolutionary dynamics; however, the maximum possible reduction is likely to be slight (Figure 3b). Additionally, it appears that mate choice is more likely to increase the variance of (realized) display traits compared to random mating when sexual selection is directional rather than stabilizing, but this may not be a general result, as we cannot explore all possible directional preference functions.

Perhaps surprisingly, in addition to increasing display trait variation, examination of the handicap model shows that sexual selection can also increase variation in loci that indicate “condition” that is associated with viability and affects the expression of male displays. Intuitively, since mate choice acts on the expressed display trait, the mechanisms above by which sexual selection increases the display trait variance also apply to any condition loci that contribute to these expressed displays.

Importantly, sexual selection via mate choice increases the variance of mating preferences, which in turn often prevents variance in the display trait from decreasing. The underlying mechanisms for this increase in preference variation are similar to those for display trait variation. First, variation of preferences among females can exaggerate display trait variance in males, and since preference and display trait in males are positively correlated, the genetic variance of preferences in males will be indirectly increased. Second, mate choice will generate assortative mating by preference phenotypes, which contributes to the between-family preference variance in the offspring generation, as described for display traits above. Since the second force dominates, sexual selection always increases the variance of preferences. As a consequence of these coevolutionary processes, variation in traits and preferences mutually promote an increase in each other, in a manner analogous to a “runaway process” but in terms of their genetic variances.

The results presented assume that a genetic correlation between traits and preferences occurs solely through non-random mating. However, the presence and magnitude of a genetic correlation can be affected by colocalization of trait and preference alleles or by pleiotropy (Butlin & Ritchie 1989, Boake 1991, Trickett & Butlin 1994, Takimoto et al. 2000) — phenomena which have been found in the context of species recognition during speciation (Wiley et al. 2012, McNiven & Moehring 2013, Xu & Shaw 2019, Ritchie & Butlin 2024). In this case, sexual selection will be more likely to increase trait variance over random mating (Figure S9), since the contribution from the covariance between mated pairs (*V*_MatedPairs_), which is proportional to trait-preference correlation, will be larger. Therefore, our results may be a minimum estimate for the effectiveness of sexual selection in increasing trait variance.

The existence of the partitions of the change in variance due to mate choice identified in this analysis (Table 1) should hold generally, regardless of the genetic control of traits or of preferences. Similarly, the fact that the correlation in traits in mated pairs will contribute to an increase variance through between-family variance should generally apply (Crow & Felsenstein 1982). However, the genetic architecture of traits and the shape of fitness and preference functions may affect, to some degree, whether the other components of variance identified in this study increase or decrease due to mate choice. The infinitesimal model used in the current study should apply well to the case when traits are controlled by many small-effect loci (Barton et al. 2017, 2023). When instead traits are controlled by a few large-effect loci, the genetic value of the offspring may not be normally distributed (Parsons & Ralph 2024) and its variance may be reduced under mate choice due to increased genetic identity between paired parents (Barton et al. 2017). We have observed that the shape of preference functions (relative/absolute vs. psychophysical) can affect the likelihood that mate choice directly increases or reduces the variance of male traits. In addition, the assumed quadratic form of viability and preference functions ensures that the trait value distribution remains normal. However, trait value distributions may evolve to be multimodal if preference functions take other forms, especially when traits are affected by few large-effect loci, as found in previous individual-based simulations and adaptive dynamic models (van Doorn et al. 2004; Weissing et al. 2011), although in this case trait variance is still increased by mate choice. For these reasons, this study may be best viewed as a “proof of concept” model (Servedio et al. 2014) explaining conceptually how variance may be affected by mate choice, rather than as generally predictive.

We hope that our analysis will prompt further studies to assess the correlations of the characteristics of sexually selected systems (e.g., preference functions, preference strengths, viability selection on preferences, viability selection on traits) with measures of genetic variation on traits and preferences. In one review of empirical studies, Prokuda and Roff (2014) found that the heritability of sexually selected traits was negatively correlated with preference strength, ostensibly contradicting what we would expect from much of our parameter space, but they did not find that the heritability differed from that of traits not under sexual selection. However, heritability may be a poor indicator of genetic variance, as genetic variance and residual variance are often positively correlated (Pomiankowski & Møller 1995).

The term “the lek paradox” suggests that the mystery of trait variation and the existence of strong preferences is deepest under a lek mating system. But we find that variation is most likely to be increased when preference is strong, costs to preferences are low, and costs to male display traits are high, as is expected on leks when many aggregated males are accessible to females (e.g., Lill 1976, Kokko 1997). Roff and Fairbairn (2014), using individual-based simulations, similarly concluded that the conditions for the evolution of strong mating preferences were likely to occur in lek mating systems because those are most likely to have females that are both very choosy and able to sample many males.

The mystery of the lek paradox has predominated our thinking in the field of sexual selection for over four decades, but its existence was never firmly established from the start. Our results suggest that there may have never really been a paradox to begin with. Sexual selection is a complex coevolutionary process, one in which intuition can rarely be trusted without formal evaluation (Servedio et al., 2014).

We hope that these findings spark additional research, both theoretical and empirical, to further clarify and reconcile research on preference and trait variation.

## Materials and Methods

We use *V*_*i*_ to denote the variance in trait *i*, and *V*_*ij*_ to denote the covariance between trait *i* and *j*. Subscripts “*m*” and “*f*” denote the male and female population, respectively, and the genetic value of a trait *i* is denoted by the subscript “*g*” as *i*_*g*_. No superscript is used for values in the juvenile population before selection, superscript “∗” denotes values after viability selection but before mate choice, and “∗∗” denotes values after mate choice. For example, 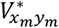 and 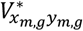 represent the phenotypic and genetic covariance between trait *x* and *y* in males after viability selection on the male trait, respectively.

We assume stabilizing viability selection on traits *z* and *x*. The relative fitness of individuals with phenotype 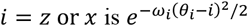. Here *θ* represents the phenotypic optimum of trait *i*, and *ω*_*i*_ is the selection strength.

We assume female preferences are subject to viability selection due to mate search. Under absolute and relative preferences, we assume that a female suffers higher costs when her most preferred male phenotype *x* or *s* is rarer, in this case further from the mean phenotype after viability selection. Under the Fisher process with absolute preferences, females with preference *y* most prefer male phenotype *x* = *y*, so their fitness is 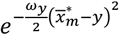, where *ω*_*y*_ is the selection strength. Similarly, with relative preferences, females with preference *y* most prefer phenotype 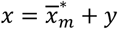, so their fitness is 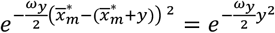. With psychophysical preferences, more choosy females (i.e., larger |*y*|) incur higher costs, so the fitness of females with preference *y* is 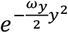(Iwasa et al. 1991; Hall et al. 2000). In the handicap model, trait *x* is replaced by the realized display trait *s* in the above fitness functions.

To derive the recursions of the genetic (co)variances, we denote the genotypic value of trait *i* within mated males and females by 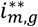 and 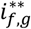, respectively. Based on the assumptions of the infinitesimal model (Barton et al. 2017), the phenotypic value of trait *i* in the offspring is

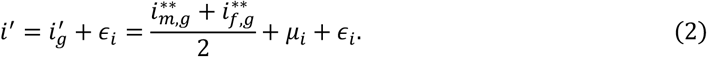

Here, *µ*_*i*_ is the deviation from the mean genetic value of the parents due to genetic processes such as segregation and recombination (Barton et al. 2017), and *ϵ*_*i*_ is environmental noise. Both *µ*_*i*_ and *ϵ*_*i*_ are normally distributed, with variances 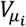 and 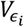, respectively. We assume that for different traits *i* ≠ *j, µ*_*i*_ and *µ*_*jj*_, as well as *ϵ*_*i*_ and *ϵ*_*jj*_, are independently distributed, so the covariance between *µ*_*i*_ and 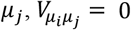, and the covariance between *ϵ*_*i*_ and 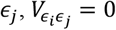.

Based on Equation (2), the genetic covariance between trait *i* and *j* after one generation is (Section 1.1 of the SI Appendix)

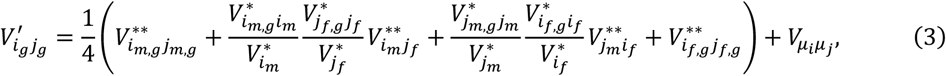

In the parenthesis of Equation (3), the first and last terms are the genetic covariance between trait *i* and *j* within mated males and females, respectively. The second and third terms represent the contribution from the covariance between paired males and females generated by mate choice, which is proportional to the covariance between the chosen male trait 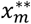 or 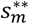 and female preference 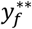 (i.e., 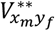 or 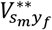 Section 1.1 of the SI Appendix). This covariance depends on the preference function. Specially, under the Fisher process, 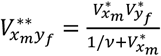 for absolute and relative preferences, and 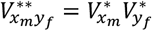 for psychophysical preferences (Fry 2024); *x* is replaced by *s* for the handicap model.

The phenotypic (co)variance between traits *i* and *j* in the next generation is 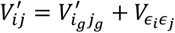 For all three preference functions, the recursions of the genetic (co)variances, and thus the equilibrium genetic (co)variances, do not depend on the mean phenotypes. However, this property may not hold for other preference functions (Figure S8).

Based on Equation (3), we can express each term in Equation (1) as

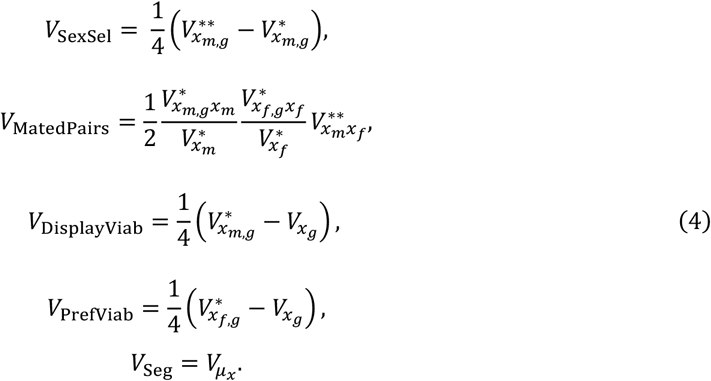

The biological meaning of each term is explained in the result section.

We track the change of phenotypic and genetic (co)variances after selection at each stage by constructing a phenotypic/genetic covariance matrix (see Section 1 in the SI Appendix). The equilibrium genetic (co)variances are obtained from numerical simulations of the recursions of genetic (co)variances (simulation code in R is available at XXX [archived upon acceptance]).

## Supplementary Information

### Section 1. Recursions of genetic (co)variances

In this section, we first describe the derivation for the recursion of genetic (co)variances. We then describe how we track the change of the phenotypic and genetic (co)variances after selection at each stage of the life cycle within a generation. In Section 1.3, we present the recursion of the genetic variance of preference *y*, which can be decomposed into several sources as we did for male trait *x* in Equation (1).

#### 1.1 Derivation for recursions of genetic (co)variances

Based on Equation (2) in the main text, the genetic covariance between traits *i* and *j* after one generation is

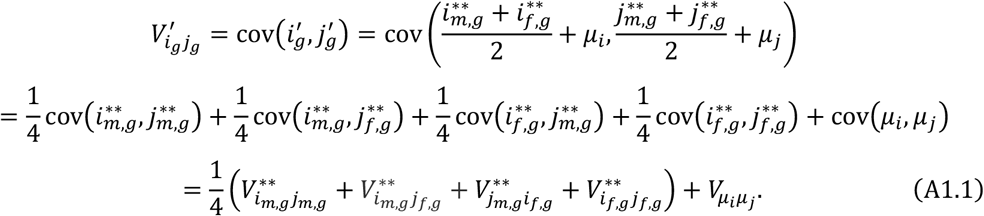

Here we derive the expression for the genetic covariance between trait *i* in mated males and trait *j* in mated females, 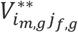 in Equation (A1.1). The term 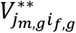 can be calculated in a similar way. By the law of total covariance, the covariance of the genetic values of trait *i* and *j* among mated pairs is

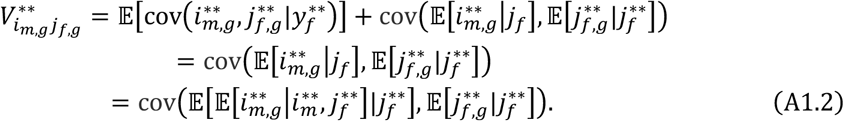

In Equation (A1.2), the term 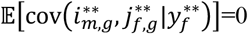 since among females with the same preference value 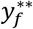, the distribution of their genetic values 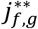 is independent of the distribution of the genetic values of trait *i* among males they mate, 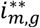. Intuitively, for females with different genetic values 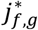, as long as they bear the same preference value 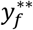, the trait value distribution among males they mate with is identical. The last expression uses the law of total expectation. Since the phenotypic and genetic values of the traits are assumed to be multivariate normally distributed, we can write Equation (A1.2) as

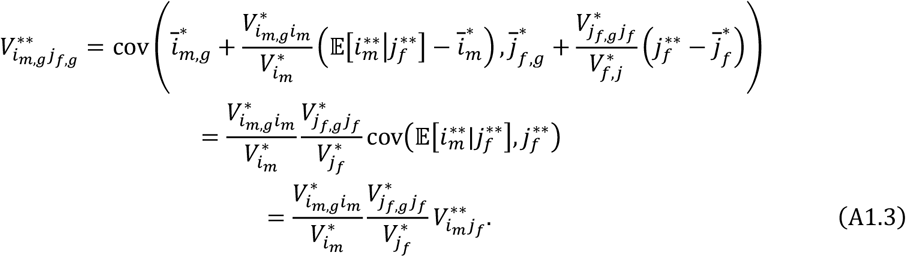

In Equation (A1.3), the first expression uses the property of conditional multivariate normal distributions. For example, since 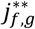 and 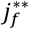 are normally distributed with covariance 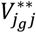, the conditional expectation is 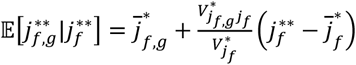. The expression for the term 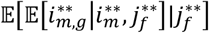 in Equation (A1.2) is written out by the same logic. The recursion of genetic covariance between trait *i* and *j* in Equation (3) is obtained by substituting Equation (A1.3) into Equation (A1.1).

Next, we calculate the phenotypic covariance between trait *i* of mated males and trait *j* of mated females, 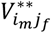 in Equation (A1.3). By the law of total covariance, we have

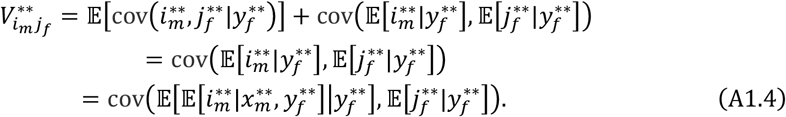

In the first expression, the term 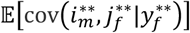 is 0 due to the same reason as explained under Equation (A1.2). The last expression uses the law of total expectation. Note that 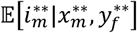 depends only on 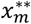 but not on 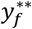. That is, based on the property of conditional multivariate normal distributions, 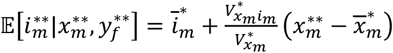. Therefore,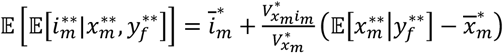 Similarly 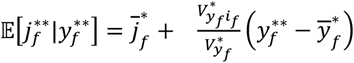. Therefore, the phenotypic covariance between trait *i* of males and trait *j* of females within mated pairs is

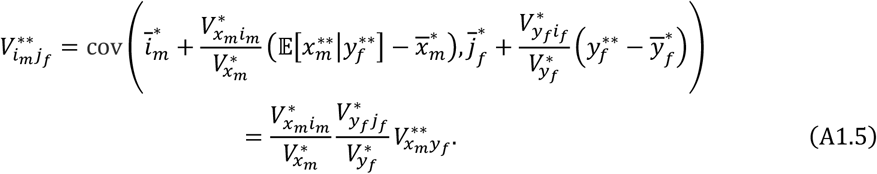

For the handicap model, where the female preference is based on the realized male trait 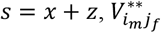 can be calculated by substituting *x* with *s* in Equation (A1.5), given by

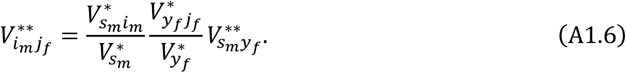

#### 1.2 Changes of phenotypic and genetic (co)variances after selection

Viability and sexual selection on a trait will alter the phenotypic variance, causing correlated changes in the phenotypic and genetic variances of other traits, as well as covariances between pairs of traits. To obtain the phenotypic and genetic (co)variances caused by selection across life stages, we consider the phenotypic/genetic covariance matrix **P** = 𝔼[***P***^T^***P***]. Here *p* is the vector of random variables, with ***p*** = *x, y, x*_*g*_, *y*_*g*_ for the model of the Fisher process and ***p*** = *x, y, z, x*_*g*_, *y*_*g*_, *z*_*g*_ for the handicap model.

Since we assume the vector ***p*** follows a multivariate normal distribution, when selection changes the variance of trait *i* from *V*_*i*_ by a fraction of *α* to (1 − *α*)*V*_*i*_, due to the properties of the conditional multivariate normal distribution, the covariance between component *j* and *k* is changed from *V*_*jk*_ to

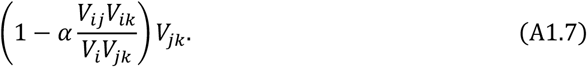

Since we assume that the fitness function of viability selection on all traits is quadratic, ***P*** remains normally distributed after selection.

For the handicap model, where mate choice is based on the realized display trait *s*, we construct a phenotypic/genetic covariance matrix after viability selection as 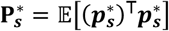, with the vector of random variables 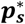 being 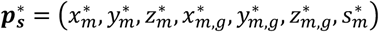. The variance of trait *s* is 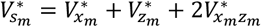, and the covariance between *s* and trait *i* ≠ *s* in 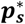 is 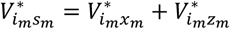.

#### 2.3 Recursion of the genetic variance of preference y

Like Equation (1) for the male trait *x*, the recursion for the genetic variance of the preference trait *y* can be decomposed into several sources as

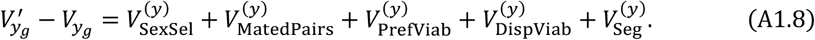

Here, 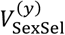 is the change that arises from the direct effect of sexual selection on male trait *x* and the genetic correlation between trait *x* and *y* within individuals. 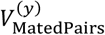 is the variance resulting from the correlation in *y* values between mated pairs under mate choice, and always causes an increase in the variance of *y* over random mating. 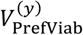 is the reduction of variance due to viability selection on preferences. 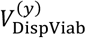 is the change in the variance of trait *y* that occurs due to viability selection on the display trait *x* in males and the genetic correlation within individuals between *y* and *x*. Mate choice, by increasing the genetic correlation between the trait and preference, may increase the magnitude of the change through *V*_DispViab_. The last term 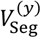 describes variance due to mutation and segregation during sexual reproduction. Based on Equation (3), the terms in Equation (A1.8) is

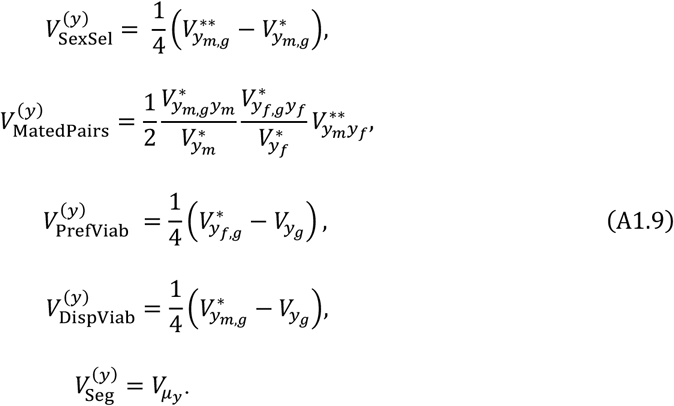

### Section 2. Effects of mate choice on variance of display trait *x* under relative/absolute preferences for the model of the Fisher process

Here we quantify the impacts of mate choice on changing the variance of trait *x* that arises from the direct effects of sexual selection (V_SexSel_ in Equation (1)), the variance resulting from correlation of trait *x* values in mated pairs (*V*_MatedPairs_), and viability selection on preference in females (*V*_PrefViab_).

Since we are interested in conditions when mate choice will increase or reduce genetic variance compared to random mating, instead of solving for the exact equilibrium variance, we focus on the effects of mate choice on changing the variance across one generation. At equilibrium, changes caused by viability selection, mate choice and the input through segregation mutation cancel each other out. In our model, the variance reduced by stabilizing viability selection is larger when the equilibrium variance is larger, so an increase (or decrease) of variance by mate choice across one generation entails larger (or lower) equilibrium variance compared to that under random mating.

Unfortunately, an analysis of how mate choice affects the genetic (co)variances at equilibrium is complicated since the expression involves heritability-like terms which will be altered after viability and sexual selection at each stage. Therefore, for simplicity and clarity, the analyses below examine the effects of mate choice under the case when there is no environmental variance, so that we can focus on the change of phenotypic variances since the heritability is always 1. Even when there is environmental variance, the analyses give qualitatively consistent results with the equilibrium genetic variances solved from recursions given by Equation (3) in *Materials and Methods* (and presented in the figures in the main text). This is because changes in the genetic variance by sexual selection ultimately occur through changes in phenotypic (co)variances.

#### 2.1 *Direct effects of sexual selection* (*V*_SexSel_)

When there is no environmental variance, by Equation (4), *V*_SexSel_ can be written as

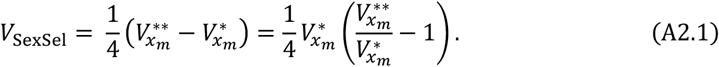

Intuitively, whether *V*_SexSel_ is negative or positive depends on whether the variance of trait *x* among males after mate choice is larger or smaller than that before mate choice (i.e., whether 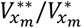 is larger or smaller than 1).

Here we show how to calculate the variance of trait *x* among males after mate choice. Denote the phenotypic distribution of the display trait *x*^∗^ in the male population after viability selection by *p*(*x*^∗^). For a female with preference *y*^∗^, the probability that she will mate with a male with display trait *x*^∗^ is

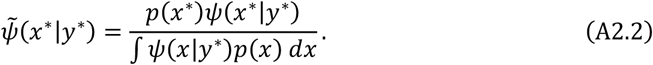

The phenotypic distribution of trait *x*^∗^ within males mated with females with preference *y*^∗^ is 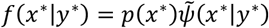. Specifically, the variance of 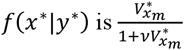 for relative and absolute preferences, and 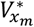 for psychophysical preferences. By integrating *f*(*x*^∗^|*y*^∗^) over the distribution of preferences in females after viability selection, *q*(*y*^∗^), we can calculate the phenotypic distribution of display trait among males after mate choice as ∫ *f*(*x*^∗^|*y*^∗^)*q*(*y*^∗^)*dy*^∗^.

Under absolute or relative preferences, the ratio of the variance of trait *x* in males before and after mate choice is

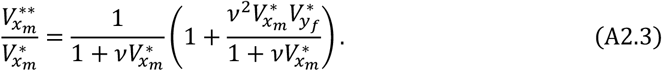

Equation (A2.3) shows that mate choice changes the variance of trait *x* among males through two effects. First, for females with the same preference *y*, the variance of trait *x* among their mates tends to differ from the variance among males before mate choice as a result of the preference function. Specifically, under relative or absolute preferences, mate choice by females with a given preference value reduces the display trait variance to 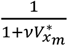 times that the variance before mating (first term in Equation (A2.3); thin blue lines in Figure 1a). This reduction increases with stronger preference strength *v*.

Second, variation in preferences among females, 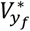, will exaggerate the display trait variation among mated males (combining across the individual blue curves in Figure 1a). This increase is captured by the second term in Equation (A2.3). The partial derivative of this term with respective to preference strength *v* is 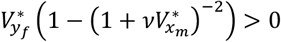, so the increase of variance through this pathway is greater with stronger preferences.

Within *V*_SexSel_, mate choice increases the variance of trait *x* (i.e., *V*_SexSel_ > 0) when the second effect outcompetes the reduction caused by the first effect. By Equation (A2.3), this requires

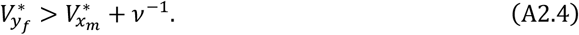

The equation shows that given 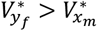, the direct effects of sexual selection through mate choice, *V*_SexSel_, increase the variance of trait *x* when preference strength *v* exceeds the critical value 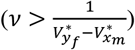. However, when 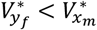, the effects of mate choice through *V*_SexSel_ always reduce the variance of trait *x* for any preference strength.

To see how changes in the preference strength affect the variance of trait *x* after mate choice 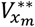, note that

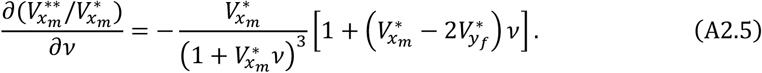

Within this pathway, stronger preference strength leads to a larger variance of trait *x* after mate choice (i.e., when Equation (A2.5) is positive) when 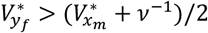, or equivalently 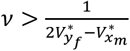. As shown before, stronger preferences raise the magnitude both of the reduction of display trait variance per preference phenotype and of the increase of display trait variance due to variation among preferences. The current result suggests that when preference strength is weak 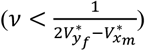, stronger preferences are more effective in raising the magnitude of the reduction through the former effect than of the increase by the latter effect, and *vice versa* when preference strength becomes strong enough 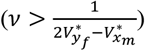.

As a summary, given fixed 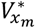 and 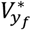, and assuming 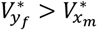, the direct effects of sexual selection through mate choice, *V*_SexSel_, will increase the variance of trait *x* when preference strength is sufficiently strong. As preference strength *v* increases from 0 (i.e., random mating), the direct effects of sexual selection *V*_SexSel_ declines from 0, reaches its minimum at

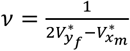, and then increases to be positive when *v* exceeds 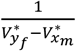. In contrast, when 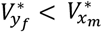, mate choice reduces the variance of trait *x* for any preference strength (*V* < 0). This reduction is greatest, and thus *V*_SexSel_ reaches the minimum value, at an intermediate preference strength 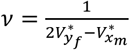 when 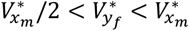, but if 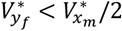, the reduction increases monotonically with stronger preference strength. These non-monotonic effects of preference strength on *V*_SexSel_ are illustrated in Figure S2.

#### 2.2 *Covariance between mated pairs* (*V*_MatedPairs_)

When there is no environmental variance, *V*_MatedPairs_ in Equation (1) can be written as

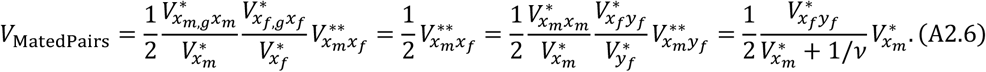

In Equation (A2.6), the expression for 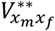 uses the result from Equation (A1.5). The term 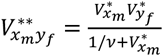 is the covariance between trait *x* of males and preference *y* of females within mated pairs, given by Table 1 in Fry (2024). Equation (A2.6) shows that *V*_MatedPairs_ is always positive and is larger as preference strength *v* increases. When mating is random, which corresponds to the case when the preference strength *v* = 0, the covariance 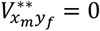 and thus *V*_MatedPairs_ = 0, as expected.

#### 2.3 *Variance change correlated with viability selection on preference* (*V*_PrefViab_)

Viability selection on preferences in females can change variance of preference, which causes a correlated change in the variance of trait x due to genetic correlation between trait *x* and *y* within individuals. Mate choice can amplify this correlated change by increasing the genetic correlation between trait *x* and *y*. Here we aim to show that the effects of mate choice on variance of male trait is small compared to the effect of *V*_MatedPairs_, and thus can be ignored.

In our model, stabilizing viability selection on preferences will reduce the variance of *y* from *V*_*y*_ to 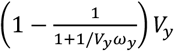. Therefore, based on Equation (A1.7), the contribution from the correlated change in the variance of trait *x* is

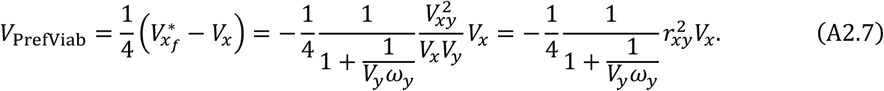

Suppose the correlation between *x* and *y* is caused solely by non-random mating through mate choice, so that *r*_*xy*_ = 0 under random mating. In this case, the change of variance caused by mate choice through *V*_PrefViab_ is proportional to the square of the correlation coefficient *r*_*xy*_. In contrast, the contribution through *V*_MatedPairs_ is proportional to 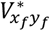 (see Equation (A2.6)), and thus is proportional to *r*_*xy*_. Therefore, the change in the variance of trait *x* due to mate choice through *V*_PrefViab_ is at higher order of *r*_*xy*_ compared to V_MatedPairs_. Therefore, the effect of mate choice through *V*_PrefViab_ tends to be much smaller than V_MatedPairs_, as illustrated in Figure S2, and thus is ignored in our analyses below.

#### 2.4 *Overall effects of mate choice* (*V*_SexSel_ + *V*_MatedPairs_)

Overall, mate choice will increase the display trait variance under two scenarios: 1) mate choice increases the variance through both *V*_SexSel_ and *V*_MatedPairs_, or 2) mate choice reduces the variance through *V*_SexSel_, but this reduction is outweighed by the increase through *V*_MatedPairs_.

The overall impacts of mate choice on the variance of display trait via the two pathways can be examined by taking the derivative of the sum of V_SexSel_ and *V*_MatedPairs_ with respect to the preference strength *v*,

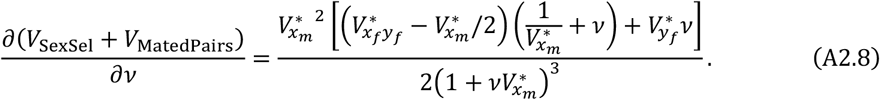

Equation (A2.8) indicates that when the covariance between the display trait and preference within choosing females is larger than half of display trait variance in chosen males 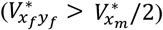, stronger preference strength will always increase the display trait variance. However, this situation will require preference variance to be sufficiently large compared to the trait variance.

When the above condition is not met 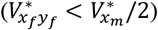, Equation (A2.8) can be rearranged to show that for a stronger preference to exaggerate display trait variance, two conditions must be satisfied: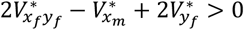, and 2) the preference strength *v* must be strong enough (see the upper-left corner in Figure 2a). The first condition can be rewritten as

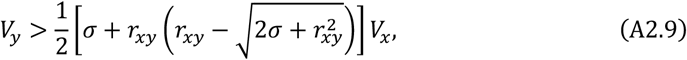

where 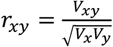 is the phenotypic correlation between the display trait and preference before selection. The parameter 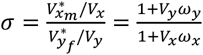 is the reduction of phenotypic variance in trait *x* compared to that in trait *y* caused by viability selection. Substituting *σ* by 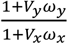 and solving for *V*_*y*_, Equation (A2.9) can be rearranged as

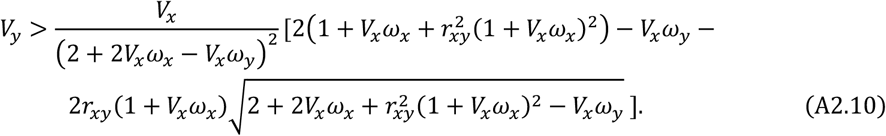

Mate choice is more likely to increase the display trait variance when the right-hand-side of Equation (A2.10) is smaller. This occurs when the correlation between the display trait and preference *r*_*xy*_ is larger, and when the strength of viability selection on the preference relative to the display trait is weaker (i.e., smaller *σ*). Based on Equation (A2.10), it can be shown that when selection on the display trait is stronger than on the preference, Equation (A2.10) can hold even when *V*_*y*_ is much smaller than *V*_*x*_.

Therefore, when the phenotypic variance of female preference is not too small and when selection on preference is sufficiently weak (small *σ*), so that Equation (A2.9) holds, the equilibrium genetic variance of trait *x* first decreases and then increases as the preference strength increases (the left-hand side in Figure 2a). However, when viability selection on the preference is strong compared to that on the male trait (large *σ*) so that Equation (A2.9) does not hold, then stronger preference consistently reduces the equilibrium genetic variance of trait *x* (left region in Figure 2d).

### Section 3. Effects of mate choice on the variance of trait *x* under relative/absolute preferences for the handicap model

Here we examine the effects of mate choice on the variance of trait *x* through the two pathways *V*_SexSel_ and *V*_MatedPairs_ under relative/absolute preferences for the handicap model. The analyses are similar to those for the model of the Fisher process in Section 2. Under the handicap model, mate choice is based on the realized display trait phenotype *s* = *x* + *z*. Since the display trait *x* and the general viability trait *z* have symmetrical contributions to trait *s*, the analyses for trait *x* can also qualitatively apply to the effects of mate choice on the variance of trait *z*. Again, for tractability, we assume there is no environmental variance, but the results qualitatively hold when environmental variance is present (see Figure 3 in the main text).

#### 3.1 *Direct effects of sexual selection* (*V*_SexSel_)

In Equation (A2.1), we have seen that the change of variance of trait *x* due to the direct effects of sexual selection is 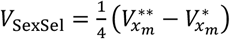. Since mate choice in the handicap model acts on the realized male trait *s*, we need to first calculate how mate choice alters the variance in trait *s*.

The phenotypic variance of the realized male trait *s* after viability selection but before mate choice is 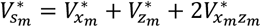. Analogous to Equation (A2.3), the phenotypic variance of *s* in males after mate choice relative to that before mate choice is

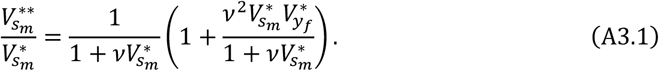

Since traits *x* and *s* are multi-normally distributed and mate choice reduces the variance of trait *s* in males by a fraction 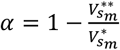, by Equation (A1.7), the variance of trait *x* among males will change from 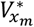 to

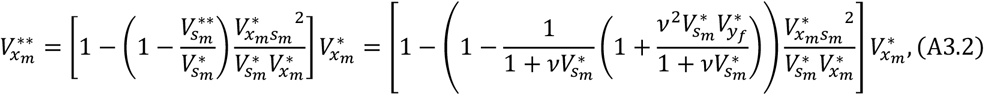

where the last expression uses the result in Equation (A3.1).

#### 3.2 Covariance between mated pairs (V_MatedPairs_)

When there is no environmental variance, the contribution from covariance in trait *x* between mated pairs is 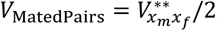. The phenotypic covariance between trait *x* in paired males and females can be obtained based on Equation (A1.6) as

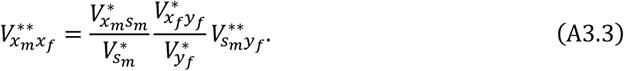

In Equation (A3.3), the phenotypic covariance between trait *s* in mated males and trait *y* in mated females is (Table 1 in Fry (2024))

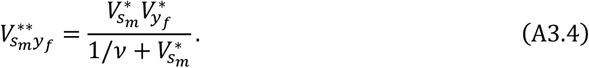

#### 3.3 *Overall effects of mate choice* (*V*_SexSel_ + *V*_MatedPairs_)

Again, we ignore the effect of mate choice on the change of variance of trait *x* through the term *V*_PrefViab_ in Equation (1), since it is much smaller than *V*_MatedPairs_ (see Section 2.3). Based on Equations (A3.2) and (A3.3), the overall effects of an increase in the preference strength *v* on the variance of trait *x* through *V*_SexSel_ and *V*_MatedPairs_ are

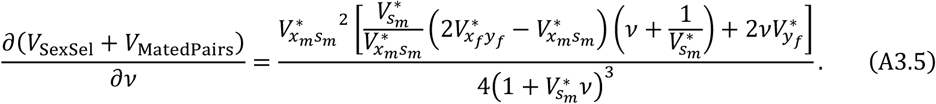

Therefore, Equation (A3.5) will be positive when the expression in the square bracket is positive. One condition for this to occur is when both terms in the square bracket is positive, which occurs when 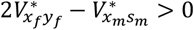 or equivalently 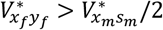.

When the first term in the square bracket in Equation (A3.5) is instead negative (i.e., when 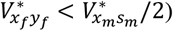, we can obtain the conditions for the expression in the square bracket to be positive by rearranging it as

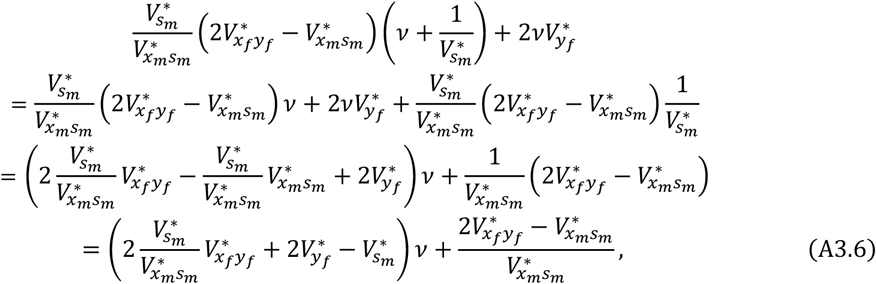

For Equation (A3.6) to be positive so that stronger preferences increase the variance of trait *x*, two conditions are required: 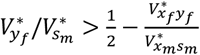, and 2) sufficiently strong preference strength *v*. Assuming for this analytical analysis that the covariance between trait *x* and general viability trait *z* is small compared to the variance of trait *x* (i.e., *V*_*xs*_ = *V*_*x*_ + *V*_*xz*_ ≈ *V*_*x*_), the first condition holds when

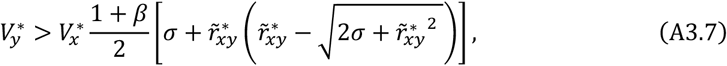

where 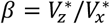 and 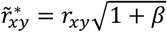. In Equation (A3.7), 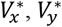 and 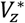 represent the genetic variances after viability selection on trait *z* but before viability selection on male. When there is no variation in general viability *z* (i.e., *β* = 0), Equation (A3.7) reduces to Equation (A2.8), derived for the Fisher process. Substituting *σ* with 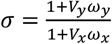, Equation (A3.7) can be rearranged as

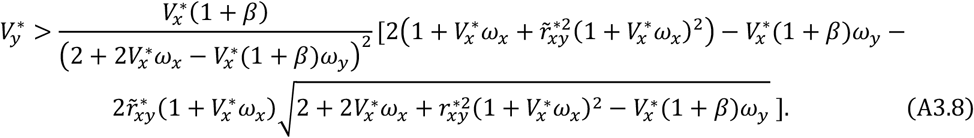

Equation (A3.8) is more likely to hold when the variance of trait *x* accounts for a larger proportion of the variance of the realized display trait *s* (i.e., smaller *β*).

### Section 4. Effects of mate choice on variance of trait *x* under psychophysical preferences

Under psychophysical preferences, ignoring the effect of mate choice through *V*_PrefViab_ (which is again likely to be small compared to the variance change through the terms *V*_SexSel_ and *V*_MatedPairs_ in Equation (1)), we find that the variance of a display trait is uniformly exaggerated by mate choice (Figures S5, S6). However, we also show in Section 4.3 that this result is not general, but due to the special function form of the psychophysical preference function.

#### 4.1 *Direct effect of sexual selection* (*V*_SexSel_)

For the model of the Fisher process, assuming no environmental variance, the change of the variance of trait *x* caused by the direct effects of sexual selection (*V*_SexSel_) is

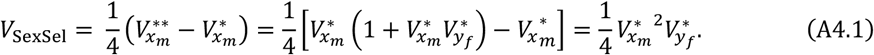

Here the variance of trait *x* after mate choice under psychophysical preferences, 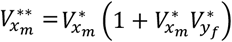, is calculated based on the method described below Equation (A2.2). Equation (A4.1) shows that sexual selection directly increases the male trait variance. On the one hand, for females with a certain preference *y*, the variance of trait *x* within the mates they mate with is the same as the variance within males before mate choice (the term 1 in 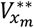). On the other hand, the term 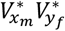 in 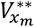 accounts for the increase in male trait variance due to variation in preference values *y* among females.

For the handicap model, under psychophysical preferences, the variance of the realized display trait *s* after mate choice compared to that before mate choice is 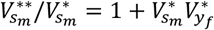 (Fry 2024). Therefore, mate choice reduces the variance of trait *s* by a fraction 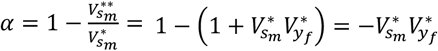. Since traits *x* and *s* are correlated, following Equation (A1.7), the variance of trait *x* after mate choice is

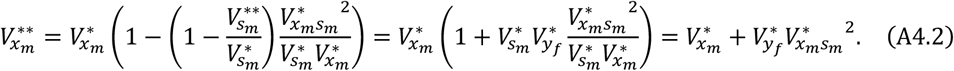

Therefore 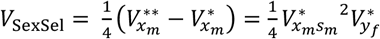.

### 4.2 *Covariance between mated pairs* (*V*_MatedPairs_)

Similar to Equation (A2.6), under the model for the Fisher process, the contribution from the covariance between mated pairs (*V*_MatedPairs_) is

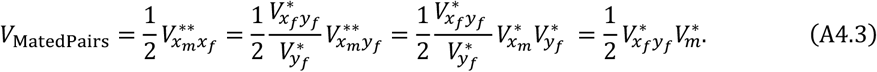

where the covariance between trait *x* of males and preference *y* of females within mated pairs, 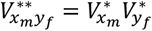, is given by Table 1 in Fry (2024). *V* is positive when there is a positive correlation between male trait *x* and preference *y* within individuals 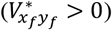. Particularly, when mating is random (i.e., when *y* = 0 for all individuals), *V*_MatedPairs_ = 0.

For the handicap model, the covariance between trait *s* of males and preference *y* of females within mated pairs is 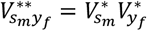. The covariance of trait *x* between mated pairs is (see Section 3.2)

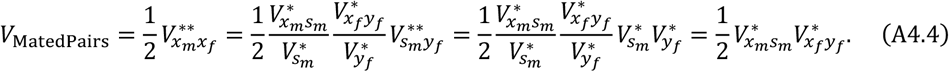

### 4.3 *Overall effects of mate choice* (*V*_SexSel_ + *V*_MatedPairs_)

Based on the results above, under psychophysical preferences the effects of mate choice on changing the variance of trait *x* through the two pathways *V*_SexSel_ and *V*_MatedPairs_ are

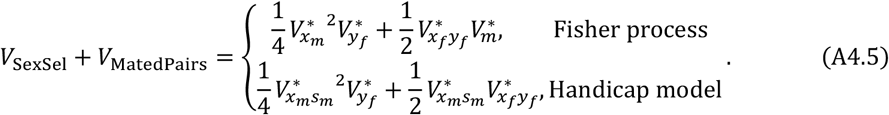

Mate choice thus always increases the variance of trait *x* over random mating through both components. However, when the variance of male trait *x* and preference *y* is too large, the recursion of the genetic variances does not converge and will explode to infinity (Figures S6, S7).

However, the above result is not general to all open-ended preferences but is due to the special form of the psychophysical preference function. For example, if a preference function is directional but takes the logistic form 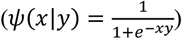, mate choice may reduce variance in the display trait when the distance between the mean phenotypic values of traits *x* and *y* is small (Figure S8), but mate choice once again always increases the preference variance (Figure S8b, S8e).

### Section 5. Effects of mate choice on the variance of preference *y* under relative/absolute preferences

Equation (A1.8) shows that mate choice contributes to the change in the variance of preference *y* through three sources. First, due to the genetic correlation between trait *x* an *y* within individuals, increases or decreases in the variance of trait *x* among males due to direct mate choice entails a correlated increase of decrease in variance in *y* in males 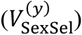. Second, mate choice generates a correlation in preference *y* values between paired females and males, which will lead to an increase in the variance of *y* over random mating, where this covariance is absent 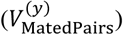. Third, mate choice increases the genetic correlation between *x* amd *y*, which amplifies the change in the variance of preference *y* correlated with the change of the variance of trait *x* caused by viability selection 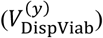.

Unlike the results for the display trait variance, we find that stronger preferences always increase the variance in preference *y*, even when sexual selection reduces the variance of the display trait (Figures 2b, 2e, 4c, 4f). This result suggests that even when the contribution of sexual selection through 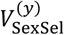 and 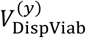 are both negative, the positive contribution through 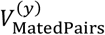 will overcompensate for them.

To see the reason for this result, under the parameter range in which the genetic variance of trait *x* is reduced by direct mate choice, the trait-preference genetic correlation, *r*_*xy*_, is weak (bottom left quadrant in Figures 2c, 2f, S4a, S4d). Additionally, as we will show below, the reduction of the preference variance through both the component 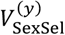 and 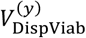 is proportional to the square of the trait-preference correlation *r*_*xy*_, while the increase through 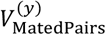 is proportional to *r*_*xy*_. Therefore, as the correlation *r*_*xy*_ approaches 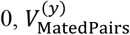 will decline in magnitude less rapidly than 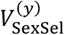, and thus tend to dominate the effect on the preference variance, as illustrated in Figure S2.

Here we show how the change of preference variance by 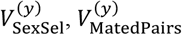 and 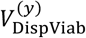 depends on the correlation coefficient between traits *x* and *y* for the model of the Fisher process. Since 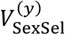 is caused by the change in the display trait variance in males due to direct mate choice (*V*_SexSel_ in Equation (1)) and the genetic correlation between *x* and *y*, we can suppose mate choice reduces the display trait variance of trait *x* among males from *V*_*x*_ to 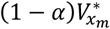. By Equation (A1.7), the correlated reduction in preference variance is

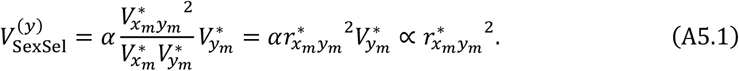

A similar analysis also applies to 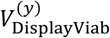. Suppose viability selection reduces the variance of trait *x* among males from *V*_*x*_ to 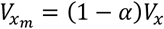. Based on Equation (A1.7), the correlated reduction in preference variance is

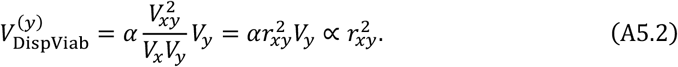

Intuitively, the correlated change in the covariance between trait *i* and *j* due to changes in the variance of trait *x* should be proportional to the production of the correlation between *x* and *i*, and the correlation between *x* and *j*. Therefore, the correlated change in the variance of *y* is proportional to 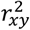.

The contribution from the correlation in preference *y* values between mated pairs is

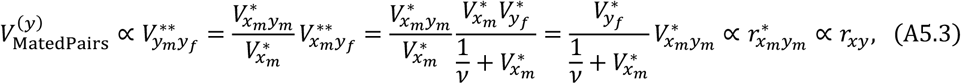

where the expression of 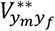 uses the result in Equation (A1.5). Intuitively, 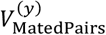 should be proportional to the covariance between display trait *x* in males and preference *y* in females, and the correlation between *x* and *y* within individual males 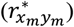. The former component depends on the mating system but not on the genetic correlation within individuals, so 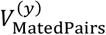 is proportional to *r*_*xy*_, instead of the square of *r*_*xy*_.

## Supplementary Figures

**Figure S1.**
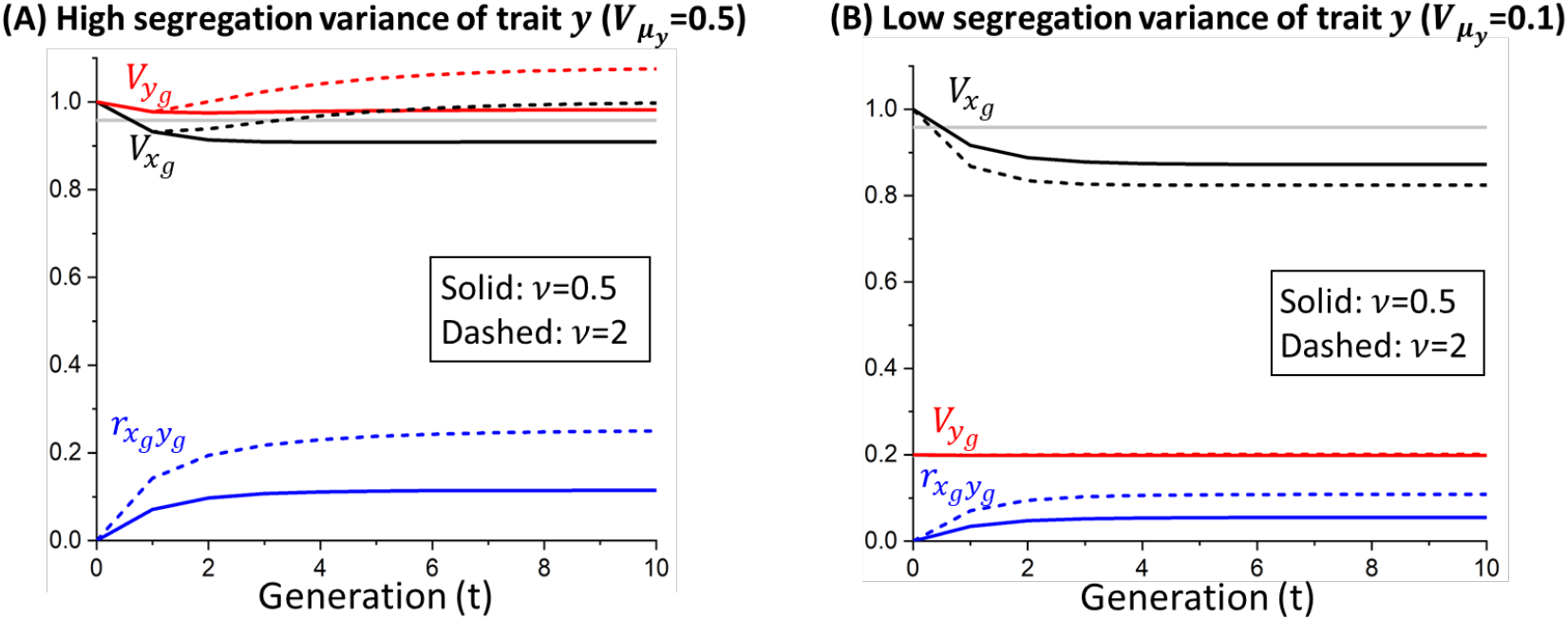
Dynamics of genetic variance and trait-preference genetic correlation at different levels of segregation variance of preference *y* and preference strength *v* under absolute or relative preferences in the model of the Fisher process. The grey line marks the equilibrium genetic variance of traits *x* and *y* under random mating. As mentioned in *Materials and Methods*, under absolute or relative preferences, the recursions of genetic (co)variances, and hence the equilibrium values, are independent of mean trait values. Results are obtained from numerical iteration of Equation (3). Parameters are 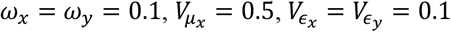.

**Figure S2.**
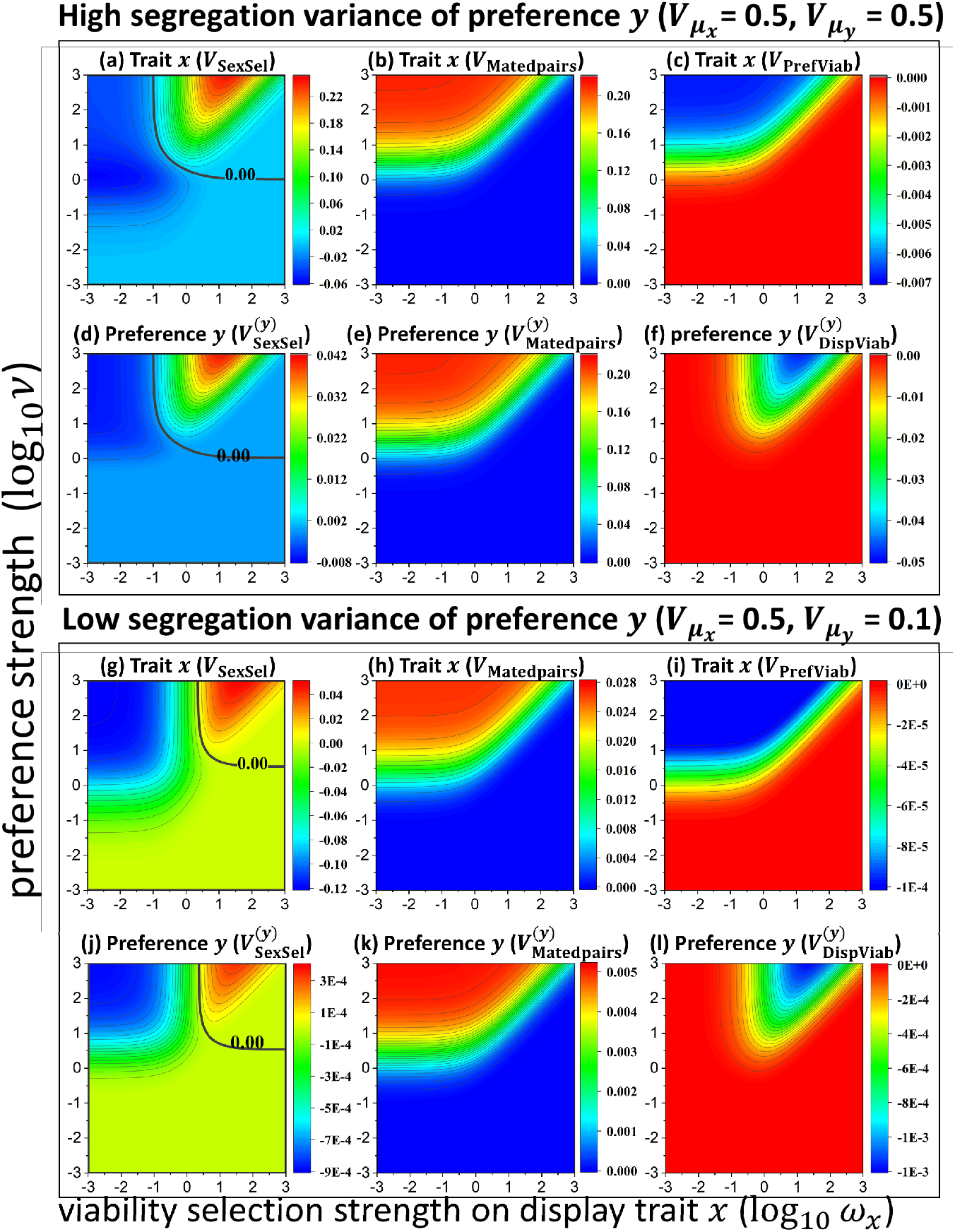
Decomposition of the effects of sexual selection on changing the genetic variance of display trait *x* and preference *y* at equilibrium for the model of the Fisher process. The changes of genetic variances by sexual selection are decomposed into three pathways *V*_SexSel_, *V*_MatedPairs_ and *V*_PrefViab_ for trait *x*, and 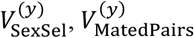 and 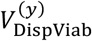 for preference *y*. The biological meaning of each pathway is detailed in Table 1 in the main text. Results are obtained from numerical simulations and the value of each component is calculated based on Equation (4) for trait *x* and Equation (A1.9) for preference *y*. Parameters are the same as in Figure 2.

**Figure S3.**
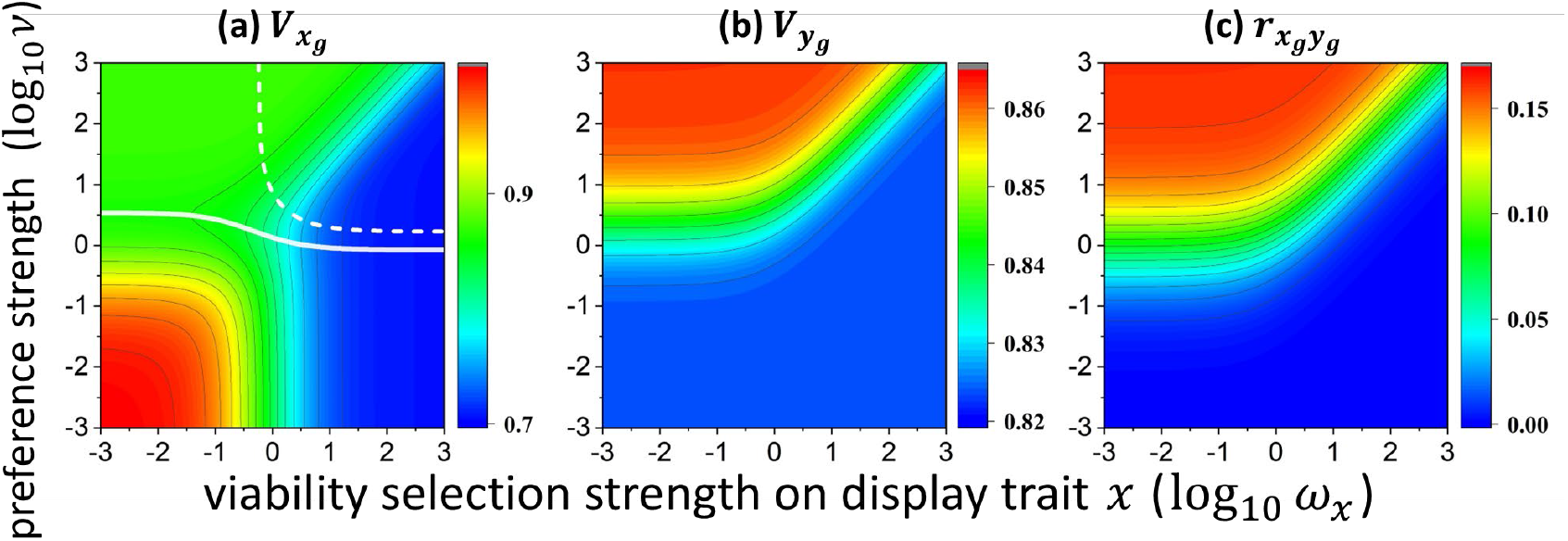
Equilibrium genetic variance of traits under relative and absolute preferences for the model of the Fisher process when viability selection on female preference is strong (*ω*_*y*_ = 1). The biological meanings of the two white lines and the values of other parameters are the same as those described in Figures 2a-c.

**Figure S4.**
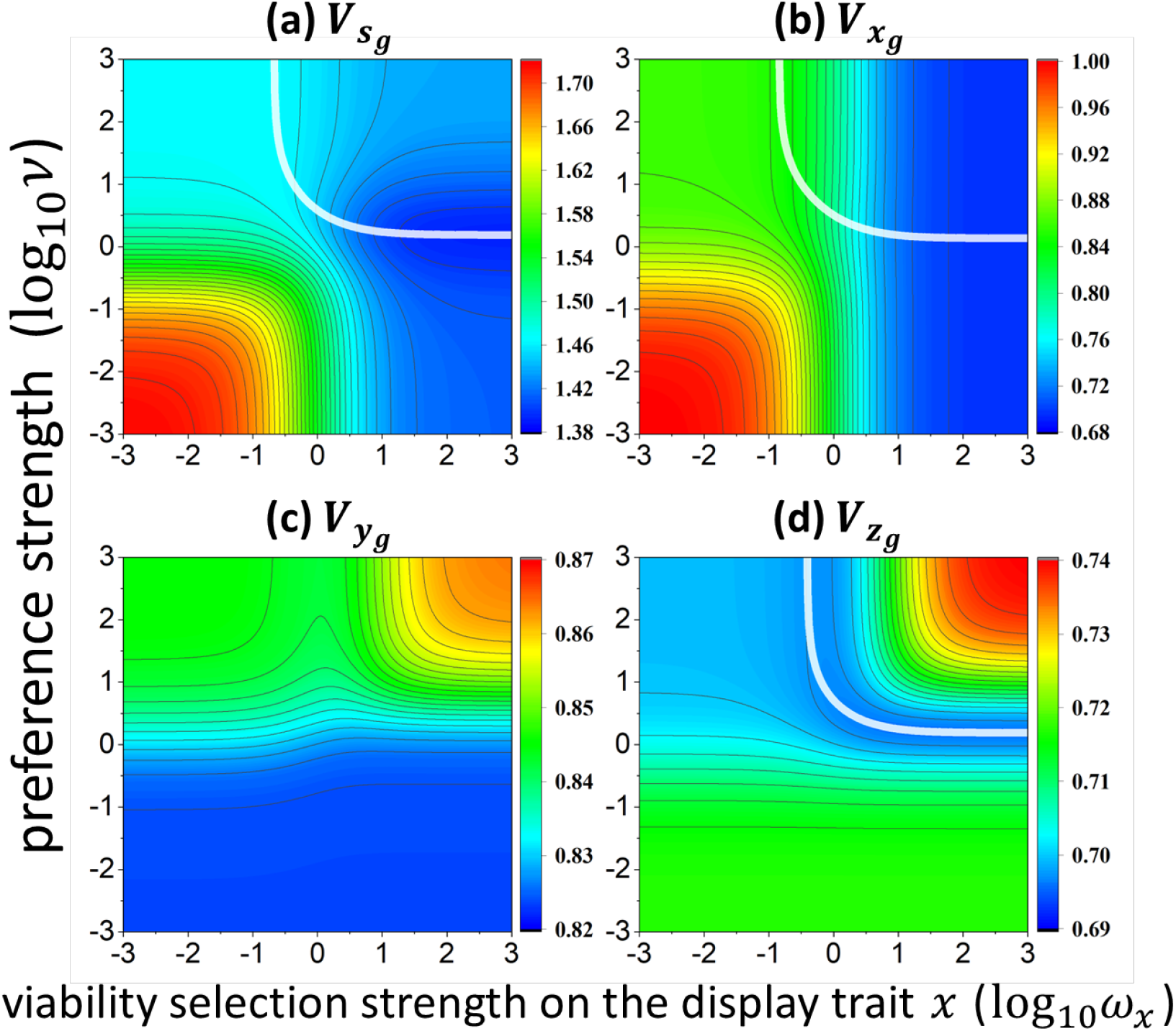
Equilibrium genetic variance of traits *s, x, y* and *z* under relative and absolute preferences for the handicap model when viability selection on female preference is strong (*ω*_*y*_ = *ω*_*z*_ = 1). The pairwise correlation between traits is very weak (not shown). Other parameters are the same as those used in Figures 4a-d. Since we set equal segregation and mutation variance for all traits, the variance of *y* is half of the variance of the realized male trait *s* when there is no selection. Under strong selection on preferences, the variation of preference in females before mate choice is quite small compared to variation of *s* unless viability selection on both *x* and *z* is very strong. Therefore, mate choice reduces the variance of trait *x* and *z* across large areas of the parameter space.

**Figure S5.**
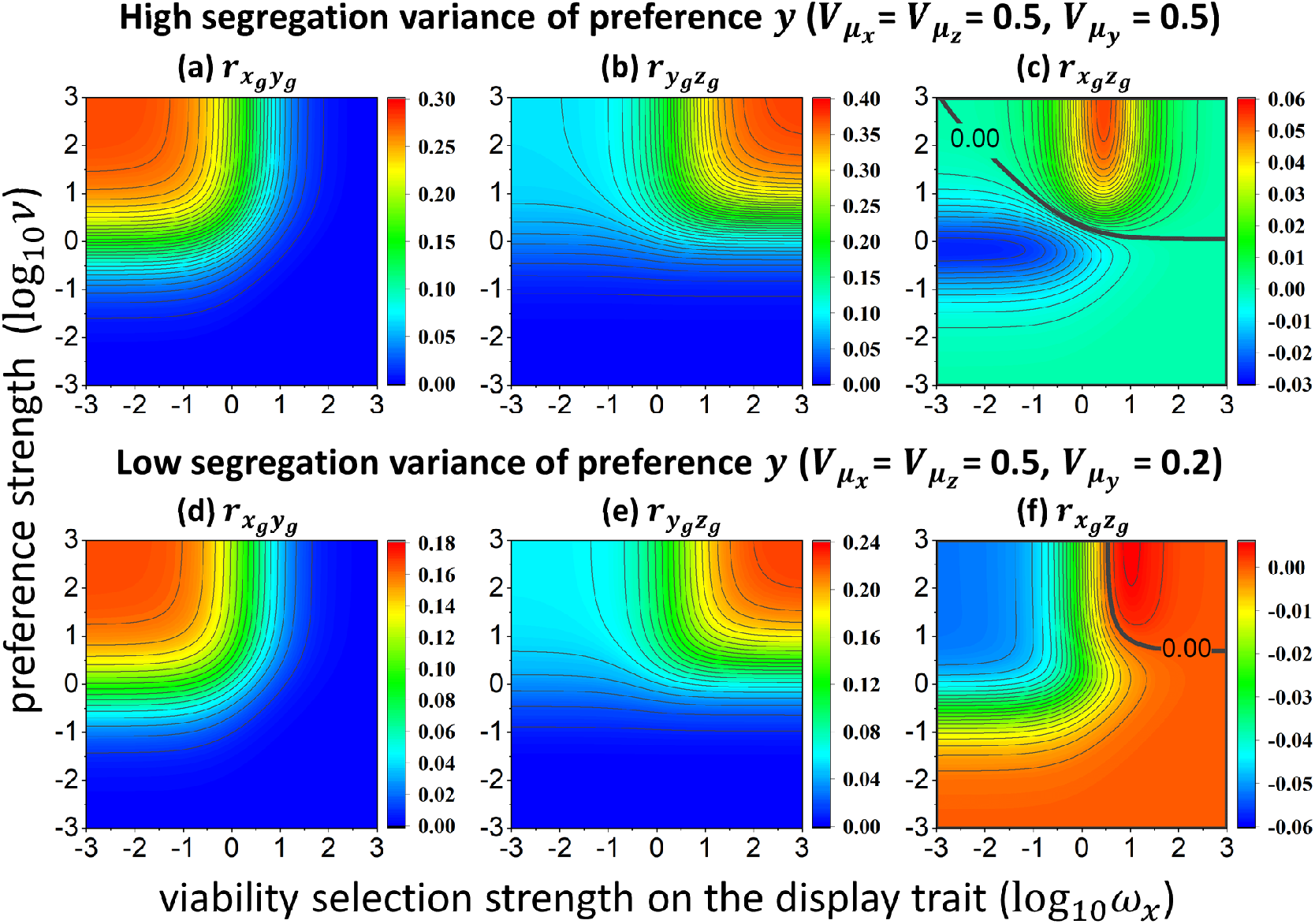
Equilibrium genetic correlation between pairs of traits (as specified for each panel) under absolute or relative preferences for the handicap model. Parameters used are the same as those used for Figure 4.

**Figure S6.**
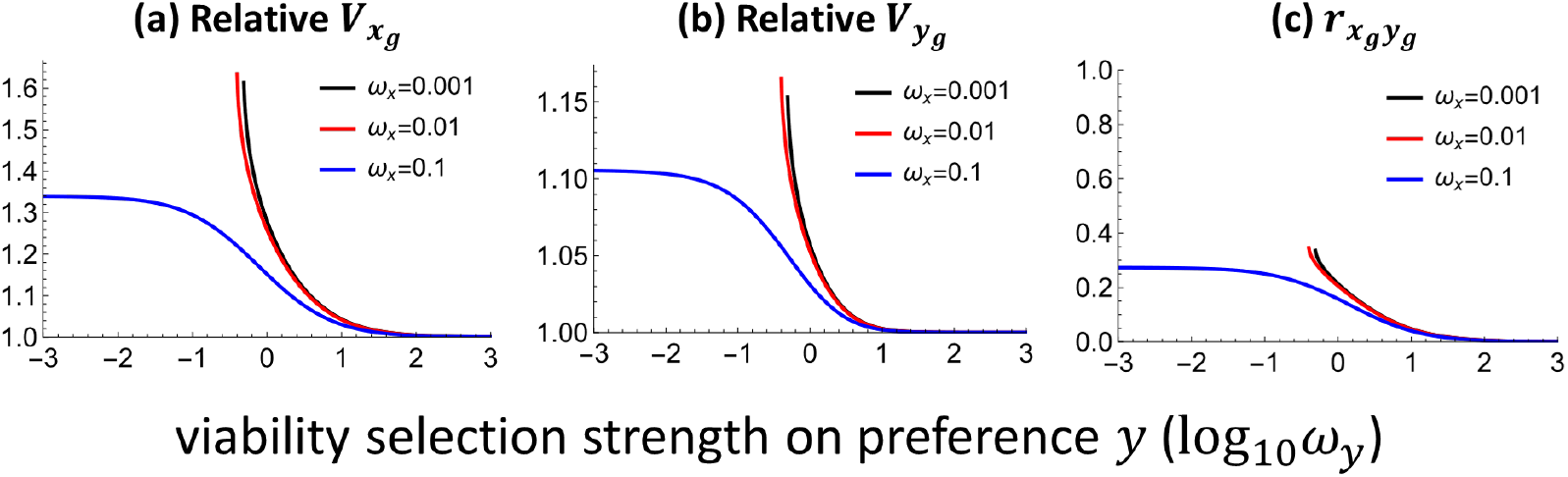
Equilibrium genetic variance and genetic correlation under psychophysical preferences for the model of the Fisher process. Panels (a) and (b) show the genetic variance of traits *x* and *y* under psychophysical preferences relative to the values under random mating. Results are obtained from numerical simulations. When viability selection strength on the preference *y* is excessively low, variation in preference *y* is large, so the genetic variances diverge towards infinity (see lines with *ω*_*x*_ = 0.001,0.01). Other parameters are 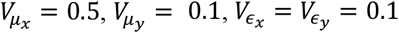

**Figure S7.**
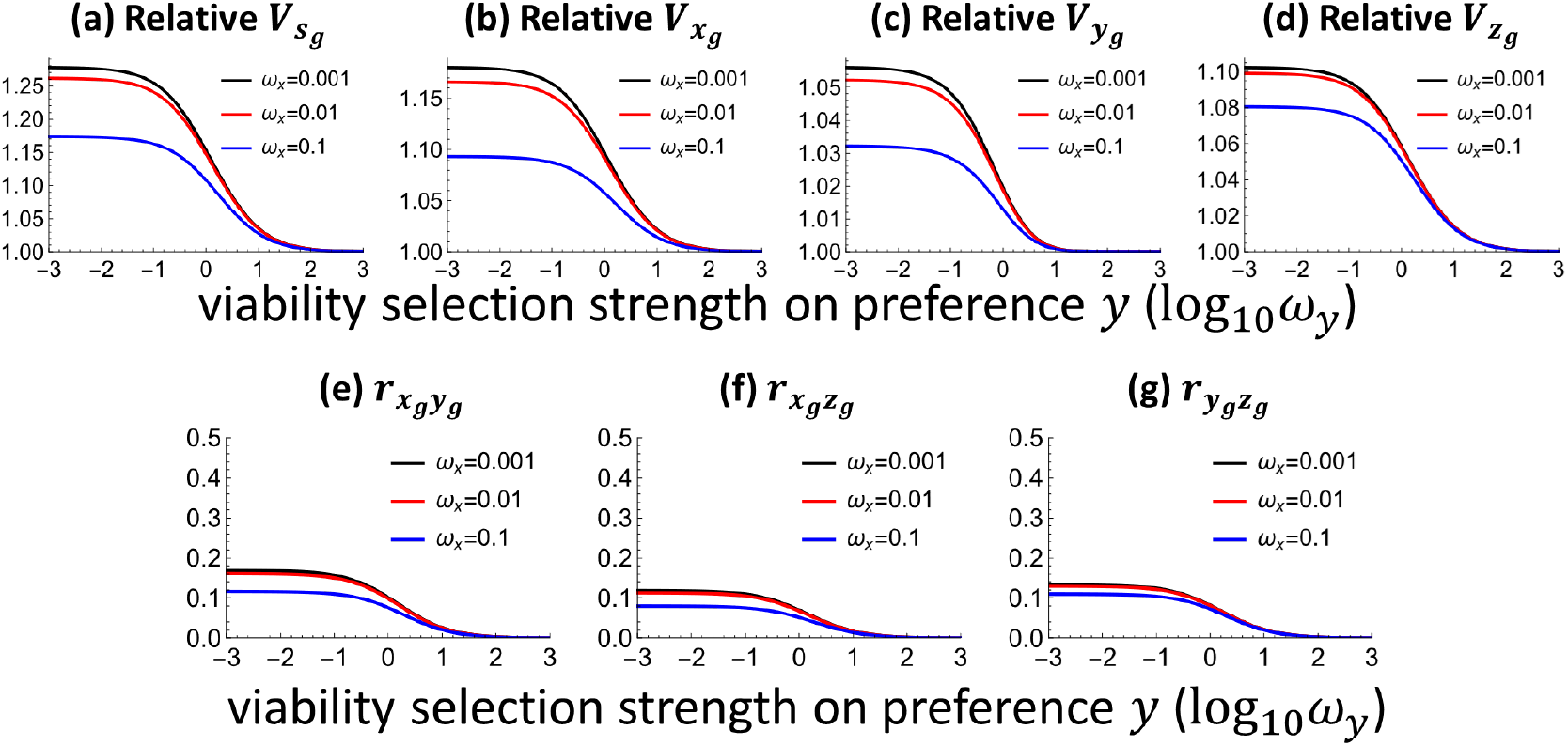
Equilibrium genetic variance and genetic correlation under psychophysical preferences for the handicap model. Panels (a)-(d) show the equilibrium genetic variance of traits *s* = *x* + *z, x, y* and *z* under mate choice relative to the values under random mating. Results are obtained from numerical simulations. The genetic variances diverge towards infinity when variation in preferences is excessively large (not presented). Other parameters are 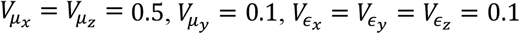.

**Figure S8.**
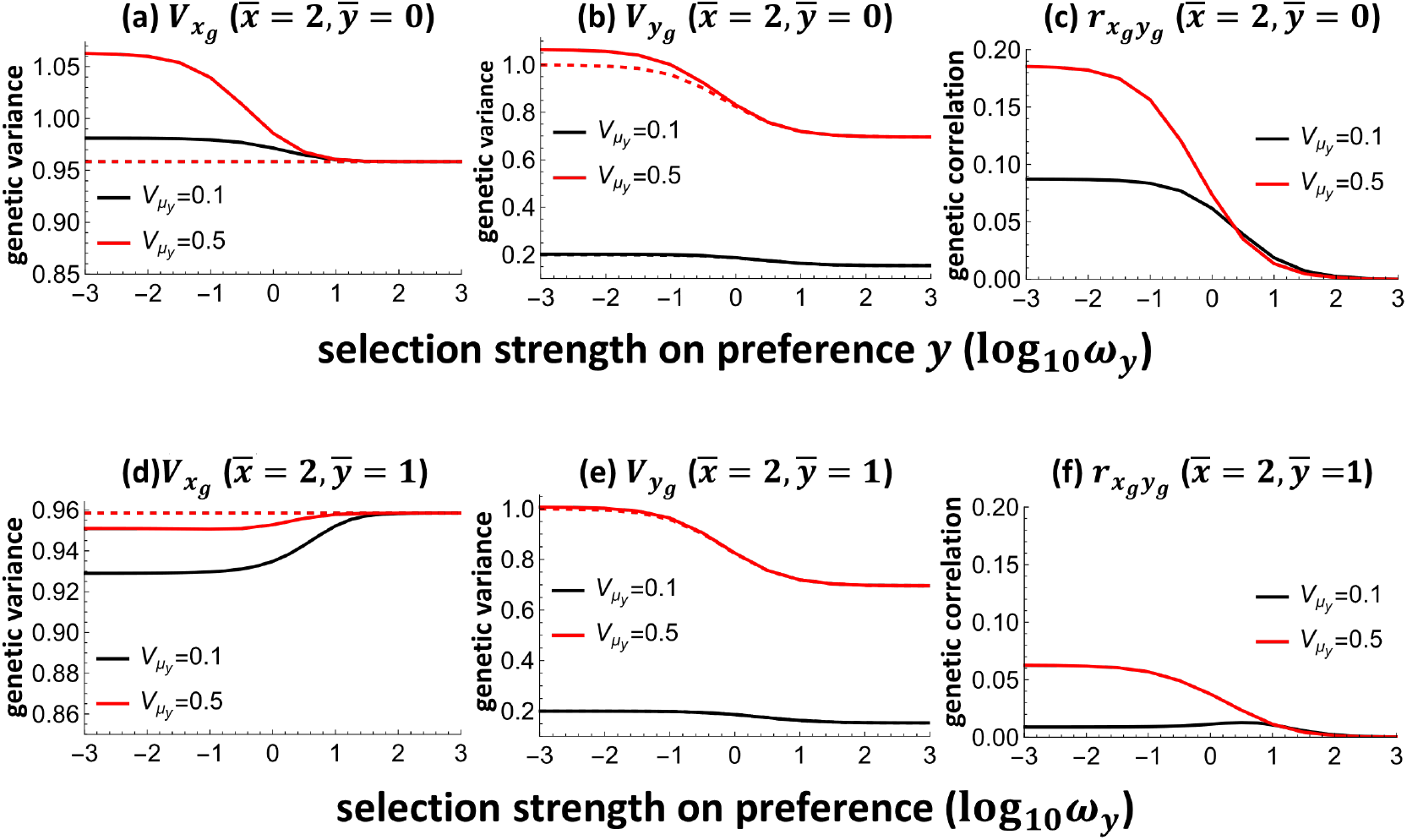
Equilibrium genetic variance of display trait *x* and preference *y*, as well as the genetic correlation between the two traits under a logistic preference function *ψ*(*x*|*y*) = 1/(1 + *e*^−*xy*^) for the model of the Fisher process. Unlike the results under psychophysical preferences, the results will depend on the mean trait values 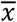 and 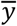, which can be seen by comparing the upper and bottom panels. Dashed lines in panels (a), (b), (d) and (e) denote the equilibrium genetic variance under random mating. Black and red lines show results under different levels of segregation variance of preference, 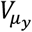. Other parameters are 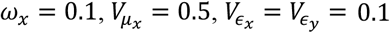.

**Figure S9.**
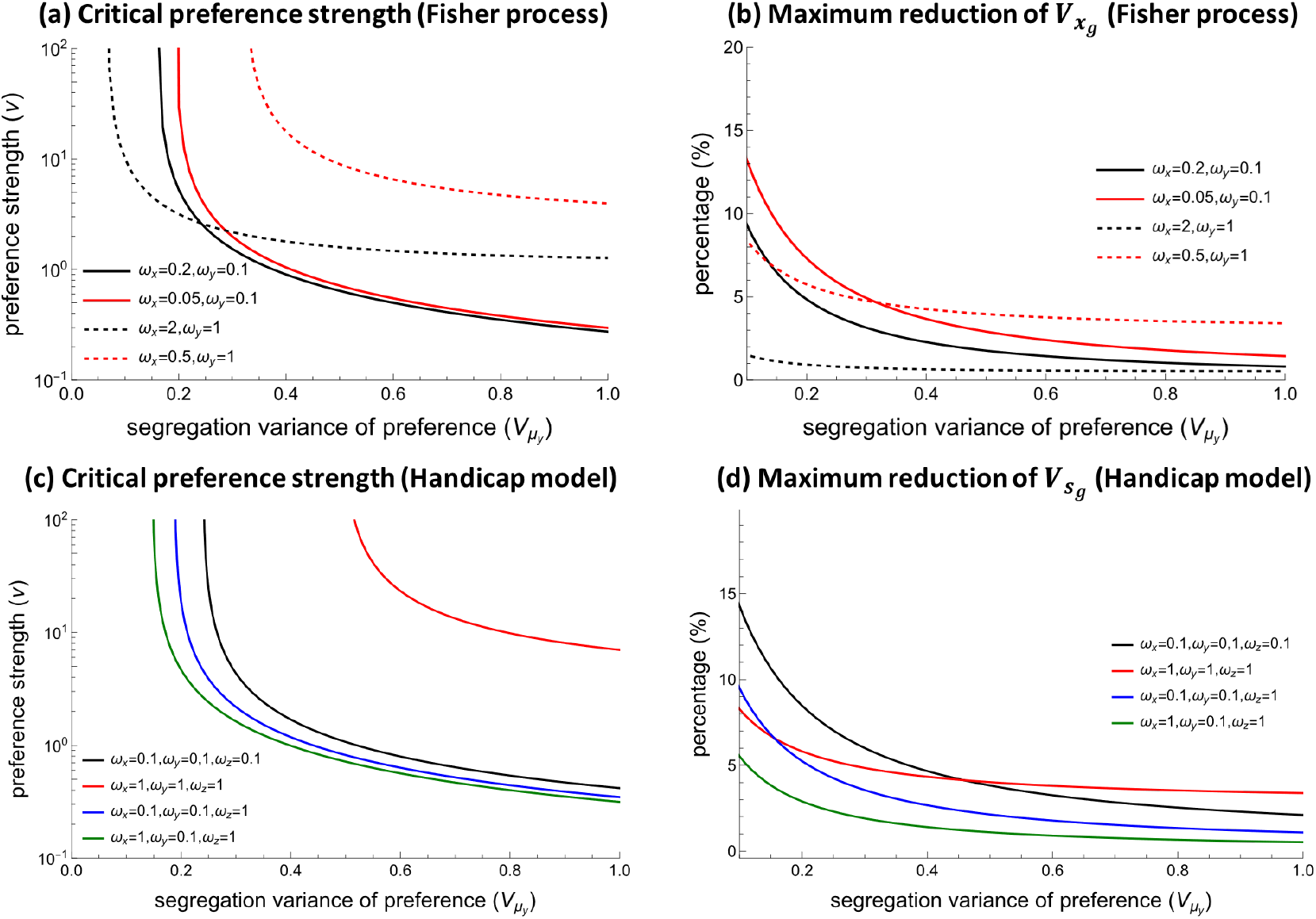
Panels (a) and (c): critical preference strength above which mate choice will lead to a larger equilibrium genetic variance of trait *x* (Fisher process) or realized display trait *s* (handicap model) over random mating under absolute or relative preferences. Panels (b) and (d): maximum possible reduction of the equilibrium genetic variance of trait *x* (Fisher process) or trait *s* (handicap model) caused by mate choice with absolute or relative preferences relative to the variance under random mating. Segregation variation in trait *x* and *y* is correlated, with correlation coefficient being 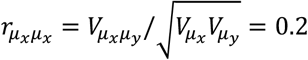. Other parameters are the same as in Figure 3. Compared to Figure 3, the conditions when sexual selection increases trait variance over random mating are relaxed and the maximum possible reduction in trait variance is reduced.

